# Novel intracellular phospholipase B from *Pseudomonas aeruginosa* with activity towards endogenous phospholipids affects biofilm assembly

**DOI:** 10.1101/2021.06.15.448513

**Authors:** Andrea J. Weiler, Olivia Spitz, Mirja Gudzuhn, Stephan N. Schott-Verdugo, Michael Kamel, Björn Thiele, Wolfgang R. Streit, Alexej Kedrov, Lutz Schmitt, Holger Gohlke, Filip Kovacic

## Abstract

*Pseudomonas aeruginosa* is a severe threat to immunocompromised patients due to its numerous virulence factors and multiresistance against antibiotics. This bacterium produces and secretes various toxins with hydrolytic activities including phospholipases A, C and D. However, the function of intracellular phospholipases for bacterial virulence has still not been established. Here we demonstrate that the hypothetical gene *pa2927* of *P. aeruginosa* encodes a novel phospholipase B named PaPlaB. PaPlaB isolated from detergent-solubilized membranes of *E. coli* rapidly degraded various GPLs including endogenous GPLs isolated from *P. aeruginosa* cells. Cellular localization studies suggest that PaPlaB is peripherally bound to the inner and outer membrane of *E. coli*, yet the active form was predominantly associated with the cytoplasmic membrane. *In vitro* activity of purified and detergent-stabilized PaPlaB increases at lower protein concentrations. The size distribution profile of PaPlaB oligomers revealed that decreasing protein concentration triggers oligomer dissociation. These results indicate that homooligomerisation regulates PaPlaB activity by a yet unknown mechanism, which might be required for preventing bacteria from self-disrupting the membrane. We demonstrated that PaPlaB is an important determinant of the biofilm lifestyle of *P. aeruginosa*, as shown by biofilm quantification assay and confocal laser scanning microscopic analysis of biofilm architecture. This novel intracellular phospholipase B with a putative virulence role contributes to our understanding of membrane GPL degrading enzymes and may provide a target for new therapeutics against *P. aeruginosa* biofilms.

## 1. Introduction

*Pseudomonas aeruginosa* causes severe hospital-associated infections, especially in immunocompromised hosts, which are complicated to treat due to the increasing antibiotic resistance and the aggressive nature of this pathogen leading to the fast progression of the infection [1, 2]. In general, the overall mortality rate determined on a large group of 213,553 patients with *P. aeruginosa* septicemia was 16 %, going along with the observation that the incidence of sepsis increases since 2001 [3]. This clearly illustrates a need for novel treatments to kill the pathogen or, at least, diminish its virulence. Therefore, WHO has recently classified *P. aeruginosa* in the group of most critical pathogens [1] and advised research and development of new antibiotics against it. Unfortunately, despite intensive investigations towards understanding virulence in this human pathogen, many genes encoding putative virulence factors remain uncharacterized [4].

In *P. aeruginosa*, as well as in other bacterial pathogens, phospholipases, the hydrolases with membrane phospholipid-degrading activity, play an important role during infections. They are classified into several groups depending on which ester bond of a glycerophospholipid (GPL) they hydrolyze [5]. While phospholipases C (PLC) and D (PLD), respectively, hydrolyze the glycerol-oriented and the head group-oriented phosphodiester bonds of phospholipids, phospholipases A1 (PLA1) and A2 (PLA2) release fatty acids bound at the *sn*-1 or *sn*-2 positions, respectively. Phospholipases B (PLB) cleave *sn*-1 and *sn*-2 bonds of GPL with similar specificity. Lysophospholipids, degradation products of PLA1 and PLA2, are converted by lysophospholipases A (lysoPLA) to glycerophosphoalcohol and fatty acid.

The contribution of bacterial phospholipases to virulence is predominantly related to damaging the host cells, which mostly enhances the survival and spread of the pathogen in the host [6, 7]. Several phospholipases of *P. aeruginosa*, namely phospholipases A ExoU [8] and PlaF [9, 10], phospholipase A/esterase EstA [11], phospholipase C PlcH [12], and two phospholipases D, PldA and PldB [13], were suggested to be virulence factors in that way. The ExoU, PldA, and PldB are directly secreted into eukaryotic cells, where they modulate native host pathways to facilitate invasion by *P. aeruginosa* or inflammation [14]. An EstA of *P. aeruginosa*, which is anchored to the outer membrane with the catalytic domain protruding into the extracellular medium, was shown to effects virulence- and resistance-related phenotypes (cell motility and biofilm formation) [11]. PlcH, one of three secreted PLCs of *P. aeruginosa*, is considered as a virulence factor because (i) it exhibits hemolytic activity; (ii) it is produced during clinical infection with *P. aeruginosa* [15], and (iii) *plcH* deletion strain of *P. aeruginosa* shows attenuated virulence in mouse burn models [16]. However, despite more than three decades of research on phospholipases, still little is known about the direct action of *P. aeruginosa* phospholipases on their membranes.

On the contrary, one of the best-studied pathogens concerning phospholipases is *Legionella pneumophila*, an intracellularly replicating Gram-negative bacterium [17]. Several phospholipases of *L. pneumophila* were proposed to have a function for establishing a proper life cycle inside a host. One of them is the major surface-associated phospholipase PlaB (LpPlaB). LpPlaB is a serine hydrolase with hemolytic activity and catalytic activity towards common bacterial phospholipids and lysophospholipids containing glycerol and choline head groups [18, 19]. However, the catalytic mechanism of LpPlaB, the mechanism of targeting to the outer membrane, structural features responsible for binding to the membrane, and its effect on the host are unknown.

Here, we expressed, purified, and characterized a homolog of LpPlaB from human pathogen *P. aeruginosa* PA01, which we named PaPlaB. Comprehensive phospholipolytic enzyme activity studies revealed that PaPlaB is a promiscuous PLB and lysoPLA, which shows strong activity towards endogenous phospholipids isolated from *P. aeruginosa*. Furthermore, we demonstrated that a *P. aeruginosa ΔplaB* deletion strain produces less biofilm with a different architecture compared to the wild-type bacterium. Thus, the PaPlaB is a novel putative virulence factor of *P. aeruginosa* PA01 belonging to the poorly understood PLB family.

## 2. Material and methods

### 2.1. Sequence analysis, and structure prediction

Amino acid sequence search and alignment were performed using BLAST and alignment tools provided by the National Center for Biotechnology Information (www.ncbi.nlm.nih.gov) [20]. The sequence alignment was visualized using BioEdit software [21]. The secondary structure was predicted with the JPred4 server using a method based on hidden Markov models [22]. TMpred server (https://embnet.vital-it.ch/software/TMPRED_form.html) was used to predict transmembrane helices with the length between 17 and 33 residues. The homology-based structural model of PaPlaB was built using the Phyre2 server [23], and the TopScore tool [24] was used to validate the structure prediction.

For further validation, protein residue-residue contacts were calculated with MetaPSICOV2 server [25] using a large sequence alignment containing 3,469 sequences. A distance violation (DV) score which indicates deviation (higher score higher deviation) between the MetaPSICOV2 predicted contacts and those found in the Phyre2 model was computed according to eq. 1. Hence, DV score below 10 indicates that a structural model has the same fold as the native structure [26].

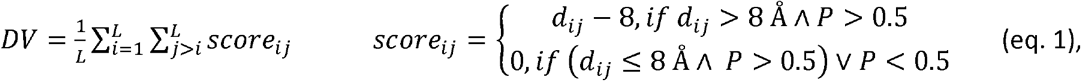

where *L* is the sequence length, *d_ij_* is the distance between C_β_ atoms of the corresponding residue pair (C_α_ if glycine is used), and *P* is the MetaPSICOV2 prediction confidence bounded between 0 and 1. The structures of oligomers were generated by *ab initio* docking of the monomeric structure followed by structural refinement using the GalaxyHOMOMER server [27].

Refined structures of PaPlaB homooligomers were used for positioning them in the membrane with PPM web server provided by Orientations of Proteins in Membranes (OPM) database [28]. The PyMol software (The PyMOL Molecular Graphics System, Version 1.2r3pre, Schrödinger, LLC.) was used for visualization of the structural model.

### 2.2. Molecular cloning

The *paplaB* gene containing the sequence that encodes a C-terminal His_6_-tag was amplified using Phusion^®^ DNA polymerase (Thermo Fisher Scientific, Darmstadt, Germany). In the PCR, the genomic DNA of *P. aeruginosa* PA01 [29], isolated with the DNeasy blood and tissue kit (QIAGEN, Germany), was used as the template together with primers paplaB_for and paplaB_rev (Table S1). The pET22-*paplaB* vector for T7 RNA polymerase-controlled expression of *paplaB* was constructed by ligation of the *paplaB* gene into the pET22b vector (Novagen, Germany) at *Nde*I and *Sac*I restriction sites, using T4 DNA ligase (Thermo Fisher Scientific). Site-directed mutagenesis of PaPlaB was performed by the Quick^®^ Change PCR method using the pET22-*paplaB* plasmid as a template and complementary mutagenic oligonucleotide pairs (Table S1) [30]. *E. coli* DH5α strain [31] was used for molecular cloning experiments. The plasmid DNA and DNA fragments from the agarose gel (1 % w/v) after electrophoresis were isolated with innuPREP Plasmid Mini Kit 2.0 and the innuPREP DOUBLEpure Kit (Analytik Jena, Germany). Oligonucleotides synthesis and plasmid DNA sequencing was performed by Eurofins Genomics (Germany).

### 2.3. Protein expression and purification

For the expression of PaPlaB with a C-terminal His_6_-tag, *E. coli* C43(DE3) [32] cells were transformed with pET22-*paplaB* plasmid, and the empty pET22b vector was used as a control. Cells were grown overnight in lysogeny broth (LB) medium [33] supplemented with ampicillin (100 μg/ml) at 37°C with agitation. Overnight cultures were used to inoculate the expression cultures to an initial OD_580nm_ = 0.05 in LB medium containing ampicillin (100 μg/ml). The cultures were grown at 37°C, and the expression of *paplaB* was induced with isopropyl-β-D-thiogalactoside (IPTG, 1 mM) at OD_580nm_ = 0.4 - 0.6 followed by incubation at 37°C for 5 hours. The cells were harvested by centrifugation (6,000 g, 4°C, 10 min) and stored at −20° C before proceeding with further analysis. Active site variants of PaPlaB carrying S79A, D196A, or H244A mutations were expressed the same as the PaPlaB.

Cells producing PaPlaB were suspended in 100 mM Tris-HCl pH 8, disrupted by a French press, and incubated for 30 min with lysozyme (2 mg/ml) and DNase (0.5 mg/ml). The cell debris and inclusion bodies were removed by centrifugation (6,000 *g*, 4°C, 10 min), and the soluble cell lysate was ultracentrifuged (180,000 *g*, 4°C, 2 h) to isolate the membrane fraction. Subsequently, the proteins were extracted from the membranes upon overnight incubation in the solubilization buffer (5 mM Tris-HCl pH 8, 300 mM NaCl; 50 mM KH_2_PO_4_; 20 mM imidazole, Triton X-100 1 % v/v) at 4°C. Insoluble debris was removed by ultracentrifugation (180,000 *g*, 4°C, 0.5 h), and the supernatant containing PaPlaB was used for purification.

Immobilized metal affinity chromatographic purification of PaPlaB was performed [34] using the ÄKTA Pure instrument (GE Healthcare). The Ni^2+^-NTA column (4 ml; Macherey-Nagel, Düren) was equilibrated with ten column volumes of the solubilization buffer before loading the sample. The column was washed with five column volumes of the washing buffer (5 mM Tris-HCl pH 8, 300 mM NaCl, 50 mM KH_2_PO_4_, 50 mM imidazole, 0.22 mM DDM) to remove unspecifically bound proteins followed by the elution of PaPlaB with 100 ml of buffer (5 mM Tris-HCl pH 8, 300 mM NaCl, 50 mM KH_2_PO_4_, 0.22 mM DDM) in which the concentration of imidazole was increased linearly from 50 to 500 mM. The fractions containing pure PaPlaB were transferred into 100 mM Tris-HCl, pH 8 supplemented with 0.22 mM DDM by gel filtration using the PD-10 column (GE Healthcare). Samples were concentrated using Amicon^®^Ultra-4 ultrafiltration device, cut-off 10 kDa (Merck Millipore). The protein was incubated at 4°C for 1 h with Bio-Beads™ SM-2 resin (Bio-Rad) equilibrated with 100 mM Tris-HCl, pH 8 to remove excess of detergent.

### 2.4. *In vitro* separation of inner and outer membranes

The separation of the inner and outer membranes of *E. coli* C43(DE3) pET22-*paplaB* (25 ml LB medium, 37°C, 5 h after induction) was performed with a continuous sucrose gradient (20 - 70 % w/v in 100 mM Tris-HCl pH 7.4). The gradients were prepared in SW40-type tubes (Beckman Coulter) using the Gradient Station (Biocomp Instruments, Canada). Isolated membranes were suspended in buffer containing 20 % (w/v) sucrose and loaded on the top of the continuous sucrose gradient followed by ultracentrifugation at 110,000 *g* for 16 h, 4°C in swinging-bucket rotor SW40 (Beckman Coulter). Fractions (1 ml) were collected from the top using the Gradient Station equipped with a TRIAX UV-Vis flow-cell spectrophotometer (Biocomp Instruments, Canada). The sucrose concentration in collected fractions was determined with a refractometer (OPTEC, Optimal Technology, Baldock UK).

### 2.5. SDS-PAGE and immunodetection

The sodium dodecyl sulfate-polyacrylamide gel electrophoresis (SDS-PAGE) was performed according to the method of Laemmli [35], and the gels were stained with Coomassie Brilliant Blue G-250. For immunodetection of PaPlaB, the gel was loaded with 10 μl of the cell, soluble and membrane fractions isolated from the cell suspension with OD_580nm_ = 25. After SDS-PAGE, proteins were transferred from the gel onto a polyvinylidene difluoride membrane [36] and detected with the anti-His (C-terminal)-HRP antibody (Thermo Fisher/Invitrogen) according to the manufacturer's instructions. The concentration of PaPlaB was determined using the UV-VIS spectrophotometer NanoDrop 2000c (Thermo Fisher Scientific). The extinction coefficient *ɛ* = 73.005 M^−1^ cm^−1^ was calculated with the ProtParam tool [37].

### 2.6. Enzyme activity assay and inhibition

Esterase activity of PaPlaB was determined in a 96-well microtiter plate (MTP) at 37°C by combining 10 μl of enzyme sample with 150 μl of the *p*-nitrophenyl butyrate (*p*-NPB) substrate [5]. Hydrolytic activities towards glycerophospholipids (GPLs) and lysoGPLs (Table S2), which were purchased from Avanti Polar lipids (Alabaster, USA), were determined by quantification of released fatty acids using NEFA assay kit (Wako Chemicals, Neuss, Germany) [5]. Lipids were dissolved in NEFA buffer (50 mM Tris, 100 mM NaCl, 1 mM CaCl_2_, 1 % (v/v) Triton X-100, pH 7.2). The enzymatic reactions were performed by combining 12.5 μl of enzyme sample with 12.5 μl of a lipid substrate (0.67 mM) at 37°C for 15 min. The fatty acid amount was calculated from the calibration curve made with 0.5, 1, 2, 3, 4, and 5 nmol oleic acid.

The inhibition of PaPlaB with PMSF, paraoxon (both were dissolved in propane-2-ol), and EDTA (dissolved in 100 mM Tris-HCl pH 8) was tested as described previously [9]. Inhibition of PaPlaB was performed by incubating enzyme aliquots with the inhibitors for 1.5 h at 30°C, followed by determination of the enzymatic activity using the *p*-NPB substrate.

Phospholipase A1 (PLA1) and phospholipase A2 (PLA2) activities of PaPlaB were measured using selective fluorescent substrates PED-A1, [N-((6-(2,4-DNP)amino)hexanoyl)-1-(BODIPY^®^FL C5)-2-hexyl-*sn*-glycero-3-phosphoethanolamine]; PC-A2_R/G_, 1-O-(6-BODIPY®558/568-aminohexyl)-2-BODIPY®FL C5-sn-glycero-3-phosphocholine (Thermo Fisher Scientific) as described previously [38]. Measurements in 96-well MTP were performed at 25°C by combining 50 μl of the substrate with 50 μl PaPlaB, or control enzymes, the PLA1 of *Thermomyces lanuginosus* (5 *U*/ml) and the PLA2 or *Naja mocambique mocambique* (0.7 *U*/ml).

### 2.7. Gas chromatography-mass spectrometric (GC-MS) analysis of fatty acids

Fatty acids were extracted after incubation of purified PaPlaB (2 ml, 4.28 μg/ml) with 1-oleoyl-2-palmitoyl-PC (PC_18:1-16:0_) (0.5 mM in 2 ml NEFA buffer) for 1 h at 37°C. After incubation 1 ml of NEFA buffer was added, and fatty acids were extracted with 12 ml CHCl_3_ : CH_3_OH = 2 : 1. The CHCl_3_ extract was removed, and FAs were extracted again with 8 ml CHCl_3_. CHCl_3_ extracts were combined and evaporated.

FAs were dissolved in 200 μl CHCl_3_. The CHCl_3_ extract was mixed with ten volumes of acetonitrile and filtered through a 0.2 μm pore size filter. The residues of the PaPlaB extracts were dissolved in 1 ml acetonitrile : methylenchloride = 4 : 1. Before GC-MS analysis, fatty acids in the PaPlaB extracts and standard solutions were derivatized to their trimethylsilylesters. For this purpose, 100 μl of each sample solution was mixed with 700 μl acetonitrile, 100 μl pyridine and 100 μl N-methyl-N-(trimethylsilyl) trifluoroacetamide and heated to 90°C for 1 h. A 1 mM fatty acid mixture in acetonitrile (C_10:0_-, C_12:0_-, C_14:0_-, C_16:0_-, C_18:0_- and C_18:1_-fatty acid (oleic acid)) was diluted to 50, 100, 200 and 400 μM and derivatized in the same manner as above. The GC-MS system consisted of an Agilent gas chromatograph 7890A and autosampler G4513A (Agilent, CA, USA) coupled to a TOF mass spectrometer JMS-T100GCV AccuTOF GCv (Jeol, Tokyo, Japan). Analytes were separated on a Zebron-5-HT Inferno column (30 m x 0.25 mm i.d., 0.25 μm film thickness, Phenomenex, USA). Helium was used as carrier gas at a constant gas flow of 1.0 ml/min. The oven temperature program employed for analysis of silylated fatty acids was as follows: 80°C; 5°C/min to 300°C, held for 1 min. The injector temperature was held at 300°C, and all injections (1 μl) were made in the split mode (1:10). The mass spectrometer was used in the electron impact (EI) mode at an ionizing voltage of 70 V and ionizing current of 300 μA. Analytes were scanned over the range m/z 50 - 750 with a spectrum recording interval of 0.4 s. The GC interface and ion chamber temperature were both kept at 250°C. After the conversion of the raw data files to the cdf-file format, data processing was performed by the use of the software XCalibur 2.0.7 (ThermoFisher Scientific). Fatty acids from the PaPlaB sample were identified by comparison of their retention times and mass spectra with those of fatty acid standards.

### 2.8. Thermal stability analysis

Differential scanning fluorimetric analysis of PaPlaB thermal stability was performed using the Prometheus NT.48 nanoDSF instrument (NanoTemper Technologies, Germany) [39]. The Prometheus NT.Plex nanoDSF Grade Standard Capillary Chip containing 10 μl PaPlaB sample per capillary was heated from 20°C to 90°C at the rate of 0.1°C/min, and the intrinsic fluorescence at wavelengths of 330 nm and 350 nm was measured. The first derivative of the ratio of fluorescence intensities at 350 nm and 330 nm as a function of temperature was used to visualize the denaturing transition and determine the “melting” temperature. Enzyme activity-based thermal stability experiments were performed by measuring the residual esterase activity of a PaPlaB sample incubated 1 h at temperatures from 30°C to 70°C [40]. After the incubation, the enzymatic assay was performed as described above using the p-NPB substrate, and the inactivation temperature was determined.

### 2.9. Multi-angle light scattering (MALS) and size-exclusion chromatography

Superdex 200 Increase 10/300 GL column (GE Healthcare) was equilibrated overnight at a flow rate of 0.6 ml/min with 100 mM Tris pH 8 containing 0.22 mM DDM. For each analysis 200 μl PaPlaB at concentrations of 1, 0.5 and 0.1 mg/ml were loaded to the column at the flow rate of 0.6 ml/min using 1260 binary pump (Agilent Technologies), and the scattered light (miniDAWN TREOS II light scatterer, Wyatt Technologies) and the refractive index (Optilab T-rEX refractometer, Wyatt Technologies) were measured. Data analysis was performed with the software ASTRA 7.1.2.5 (Wyatt Technologies) under the assumption that dn/dc of DDM is 0.1435 ml/g and the extinction coefficient of PaPlaB is 1.450 ml/(mg*cm) [41].

Size-exclusion chromatographic (SEC) analysis of PaPlaB in Tris-HCl (100 mM, pH 8, 0.22 mM DDM) buffer was performed using Biosep-SEC-S3000 column (Phenomenex, Aschaffenburg, Germany), LC-10Ai isocratic pump (Shimadzu, Duisburg, Germany) and SPD-M20A photodiode array detector (Shimadzu, Duisburg, Germany). The molecular weight (M_w_) of standard proteins dissolved in the same buffer as PaPlaB was determined (Table S3). For the analysis, 100 μl of PaPlaB or protein standard sample was loaded on the column, and separation was achieved at a flow rate of 0.5 ml/min and 26°C.

### 2.10. Construction of *P. aeruginosa ∆plaB* strain

The *P. aeruginosa ∆plaB* mutant strain was generated by homologous recombination [42]. In short, *P. aeruginosa* PAO1 cells were conjugated with the pEMG-*∆plaB* mutagenesis vector containing the 814 bp fragment of the upstream region of *paplaB*, followed by a gentamicin resistance gene and the 584 bp downstream region of *paplaB*. For that, *E. coli* S17-1 ⍰pir transformed with the pEMG-*∆plaB* plasmid was used as a donor strain. *Pseudomonas* cells with pEMG-*∆plaB* plasmid integrated on the chromosome were selected on LB-agar plates containing gentamicin (30 μg/ml), kanamycin (300 μg/ml; a kanamycin resistance gene is encoded on pEMG plasmid) and Irgasan (25 μg/ml; used for negative selection of *E. coli*). Cells transformed with the plasmid pSW-2 containing the I-SceI restriction endonuclease were cultivated on LB agar plates containing benzoic acid (2mM; for induction of I-SceI expression) and Irgasan (25 μg/ml). The deletion of the *paplaB* gene was confirmed by PCR amplification using the genomic DNA of *P. aeruginosa ∆plaB* as the template (Fig. S1).

### 2.11. Fluorescence imaging of biofilm in flow chambers

*P. aeruginosa* PAO1 and *∆plaB* biofilms were grown on a microscope cover glass (24 mm x 50 mm, thickness 0.17 mm, Carl Roth GmbH & Co. KG, Karlsruhe, Germany), which was fixed with PRESIDENT The Original light body silicon (Coltène/Whaledent AG, Altstätten, Switzerland) on the upper side of the three-channel flow chambers [43]. The flow chambers and tubes (standard tubing, ID 0.8 mm, 1/16” and Tygon Standard R-3607, ID 1.02 mm; Cole-Parmer GmbH, Wertheim, Germany) were sterilized by flushing with sterile chlorine dioxide spray (Crystel TITANIUM, Tristel Solutions Ltd., Snailwell, Cambridgeshire, United Kingdom). Afterward, the flow chambers were filled with 1 % (v/v) sodium hypochlorite, and the tubes were autoclaved. All biofilm experiments were performed at 37°C with a ten-fold diluted LB medium. Before inoculation, the flow chamber was flushed with 1:10 diluted LB medium for 30 minutes with a flow rate of 100 μl/min using the IPC12 High Precision Multichannel Dispenser (Cole-Parmer GmbH, Wertheim, Germany). For inoculation, an overnight culture of *P. aeruginosa* PAO1 or *∆plaB* was adjusted to an OD_580nm_ of 0.5 in 1:10 diluted LB medium. The diluted culture (300 μl) was inoculated in each channel. After the interruption of medium supply for 1 h, the flow (50 μl/min) was resumed, and the biofilm structure was analyzed after 24, 72, and 144 h grown at 37°C. For visualization, the cells were stained with propidium iodide and SYTO 9 dyes using the LIVE/DEAD™ BacLight™ Bacterial Viability Kit (Thermo Fisher Scientific). Imaging of biofilm was performed using the confocal laser scanning microscope (CLSM) Axio Observer.Z1/7 LSM 800 with airyscan (Carl Zeiss Microscopy GmbH, Germany) with the objective C-Apochromat 63x/1.20W Korr UV VisIR. The microscope settings for the different fluorescent dyes are shown in Table S4. The analysis of the CLSM images and three-dimensional reconstructions were done with the ZEN software (version 2.3, Carl Zeiss Microscopy GmbH, Germany). Experiments were repeated two times, each with one biological replicate that was analyzed at three different points by imaging a section of 100 x 100 μm.

### 2.12. Crystal violet biofilm assay

*P. aeruginosa* wild-type and *∆plaB* cultures incubated in LB medium overnight at 37°C in Erlenmeyer flasks (agitation at 150 rpm) were used to inoculate 100 μl culture with OD_580nm_ 0.1 in plastic 96-well MTP. Cultures were grown at 37°C without agitation, and the cells attached to the surface of MTP after removing the planktonic cells were stained with 0.1 % (w/v) crystal violet solution for 15 min, solubilized with acetic acid (30 % v/v) and quantified spectrophotometrically [44].

## 3. Results

### 3.1. *P. aeruginosa* gene *pa2947* encodes a putative phospholipase A

We performed a BLAST search using sequences of the currently known bacterial PLAs to identify novel phospholipases A (PLA) in *P. aeruginosa* PA01 [6]. One of the results was a gene *pa2927* encoding a 49.5 kDa protein with moderate sequence similarity (39 %) to LpPlaB, a major intracellular PLA of *L. pneumophila* [18, 19, 45–48]. We named the *P. aeruginosa* homolog analogously, PaPlaB. A BLAST search revealed substantial sequence identity (27 to 43 %) between PaPlaB and several uncharacterized proteins and putative phospholipases (Figs. 1a and S2). The recently annotated active site of LpPlaB (Ser85, Asp203, His251) [18] allowed us to identify Ser79, Asp196, and His244 as residues of the putative catalytic triad in PaPlaB, which are strongly conserved among PaPlaB homologs (Fig. 1a). PaPlaB, similar to LpPlaB, contains the catalytic Ser79 embedded into a VHSTG pentapeptide, which differs from the canonical GXSXG lipase motif by the absence of the glycine at position −2 of the serine. The present threonine residue was shown to be functionally important, as the mutation T83G reduced LpPlaB activity by 95 % [18]. The difference in the catalytic pentapeptide was proposed to be a novel feature of the lipolytic family represented by LpPlaB [18]. Although no conserved domains encompassing the catalytic residues were identified in PaPlaB using either NCBI’s conserved domain [49] or Pfam [50] databases, we observed a notably higher sequence identity of the 270 amino-terminal residues of PaPlaB to its homologs than the carboxy-terminal residues (271 – 443). Even secondary structure elements predicted in the N-terminal regions of PaPlaB and LpPlaB agreed better than secondary structure elements predicted in the C-terminal regions (Fig. S3).

**Fig. 1:**
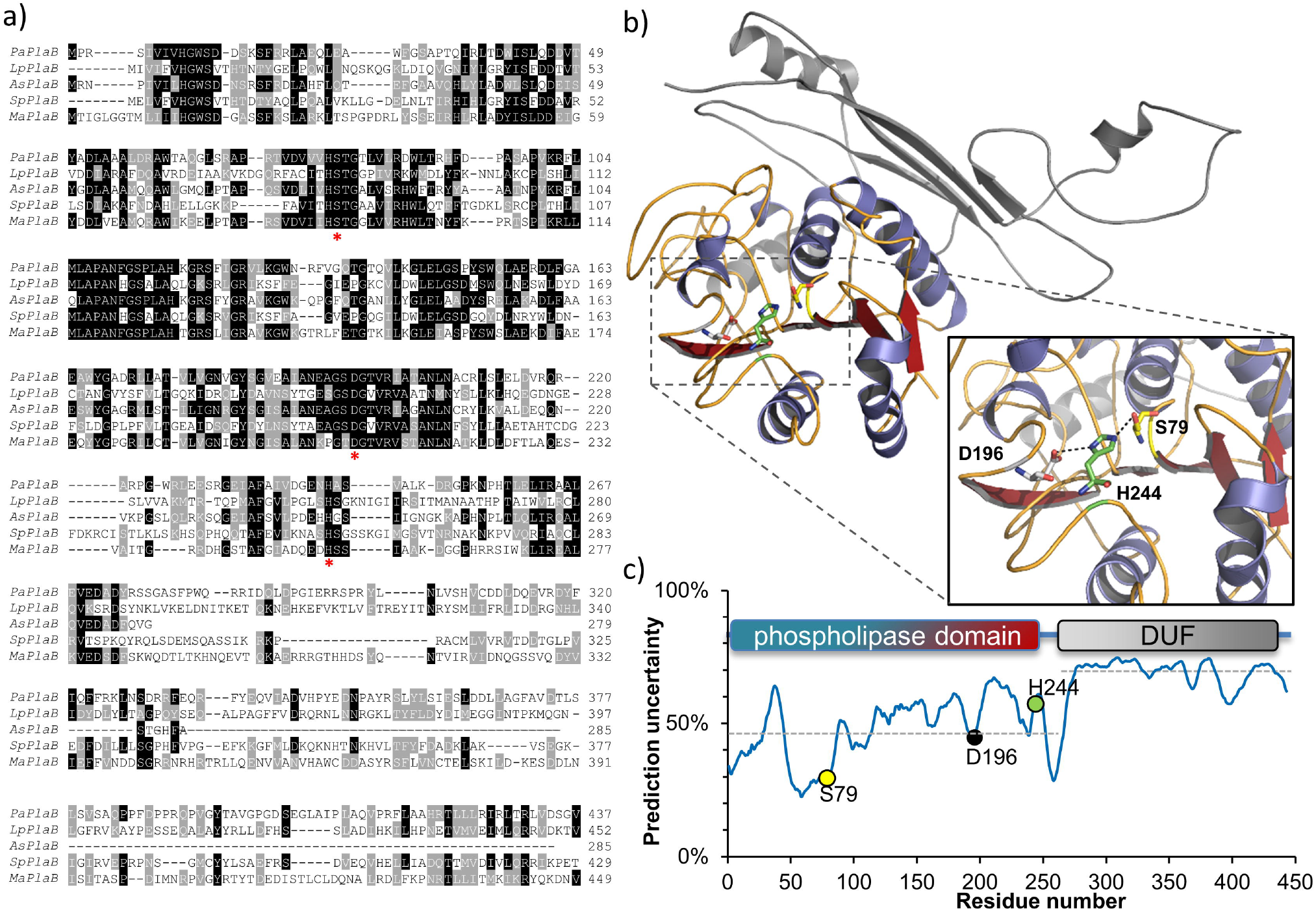
Putative two-domain architecture of *P. aeruginosa* PaPlaB. **a)** Sequence alignment of PaPlaB from *P. aeruginosa* (PaPlaB), *Legionella pneumophila* (LpPlaB), *Alishewanella sp.32-51-5* (AsPlaB), *Shewanella pealeana* (SpPlaB), and *Marinobacter algicola* (MaPlaB). Black and grey background indicate identical and similar residues in at least three proteins, respectively. The catalytic triad residues (Ser79, Asp196, and His244 in PaPlaB) are marked with asterisks. The last six PaPlaB residues are shown in the Fig. S2. **b)** PaPlaB structural model. The N-terminal domain model (coloured secondary structure elements) was based on the structure of the protein of unknown function from *L. innocua* (PDB code: 3DS8). The C-terminal domain (grey) was modelled *ab initio* using the Phyre2 web server [51]. The view of the catalytic triad is expanded. **c)** The residue-wise uncertainty of the predicted PaPlaB model was computed with TopScore, a quality assessment tool for protein structure models [24]. Grey lines indicate average uncertainties of the phospholipase and DUF domains.

To assess whether PaPlaB is a two-domain protein, we predicted its 3D structure with the Phyre2 homology modeling server [51]. Using the structure of a protein of unknown function from *Listera innocua* (PDB code: 3DS8) as a template, the homology model of the N-terminal PaPlaB domain revealed an α/β-hydrolase fold, which provides a scaffold for the canonical serine-hydrolase catalytic triad (Fig. 1b) as found in many lipolytic enzymes [52, 53]. No homolog with a resolved structure was found for the C-terminal domain of PaPlaB. Therefore, the *ab initio* approach of the Phyre2 server was used. The resulting model for this domain suggested a four-stranded β-sheet (Fig. 1b) which was furthermore validated. Validation with TopScore [24] confirmed a higher quality of the N-terminal domain than the C-terminal one (Fig. 1c). Additional validation with a MetaPSICOV2 server shows two distinct groups of residue-residue contacts, one in the N-terminal and one in the C-terminal region, thus corroborating a two-domain protein structure (Fig. S4). DV score, a measure of the deviation of predicted contacts versus those derived from the structural model, of 2.7 suggests that the PaPlaB model has a fold similar to the native structure, as DV < 10 was calculated for real structures. In summary, our sequence, secondary and tertiary structure analyses indicate that PaPlaB has an N-terminal putative phospholipase A domain with an α/β-fold and a C-terminal domain, the function of which is currently unknown.

### 3.2. Expression of *paplaB* in *E. coli* yields a membrane-bound phospholipase A

To experimentally test the putative PLA function, we set out to heterologously express and purify PaPlaB. To achieve this, we constructed the PaPlaB expression vector (pET22-*paplaB*) suitable for heterologous expression in *E. coli* strains that contain the T7 RNA polymerase gene. *PaplaB* gene in pET22-*paplaB* plasmid was modified by including a sequence coding for six histidine residues at the 3’ end to enable purification of the protein using immobilized metal affinity chromatography (IMAC) and the protein expression was conducted in *E. coli* C43(DE3) cells. SDS-PAGE (Fig. S5) and Western blot (Fig. 2) analyses of cells sampled during the first 5 h after induction revealed the expression of a protein with an estimated molecular weight (M_w_) of ~50 kDa, which agrees with the theoretical M_w_ of PaPlaB (49.5 kDa). The expression of PaPlaB variants with mutated putative catalytic triad residues S79, D196, and H244 yielded also ~50 kDa proteins as shown by SDS-PAGE (Fig. S6a) and Western blot (Fig. S6b) analyses. Esterase activities of cell lysates revealed that the wild type PaPlaB was active while all three variants showed activities comparable to the activity of the empty vector control (Fig S6c)

**Fig. 2:**
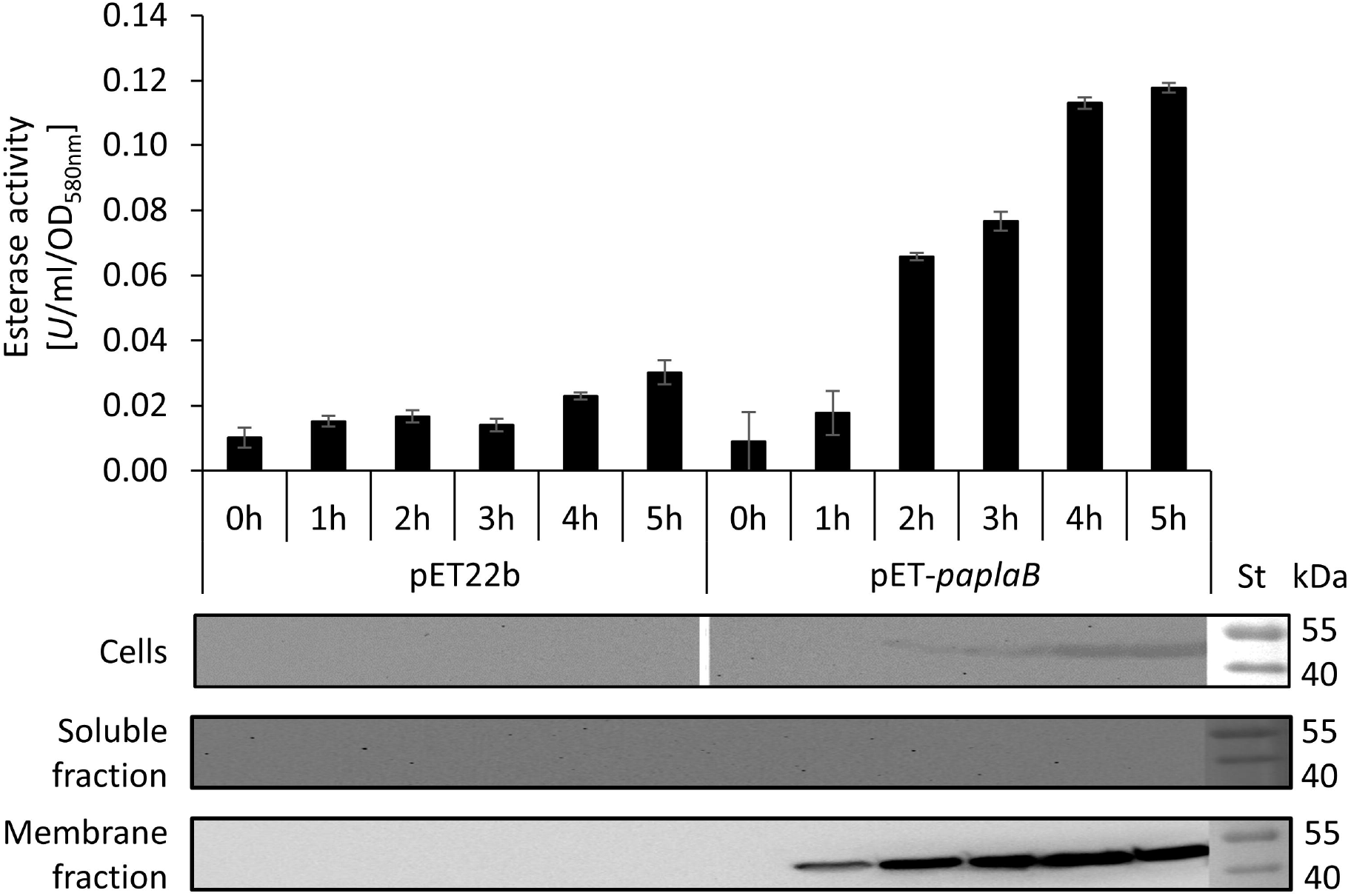
PaPlaB is heterologously expressed in *E. coli*. Expression, localization, and activity of PaPlaB was tested at 1, 2, 3, 4, and 5 h after induction. Cell lysates (10 μl, OD_580nm_ = 10) were analyzed by esterase *p*-NPB assay (top) and Western blotting against the His_6_-tag (below). Disrupted cells were fractionated by ultracentrifugation into soluble (cytoplasmic and periplasmic proteins) and membrane protein fractions that were analyzed by Western blotting. *E. coli* C43(DE3) carrying empty vector pET22b were grown under the same conditions and were used as the negative control. Molecular weights of standard proteins (St) are indicated on the right-hand side. The esterase activity results are means ± S.D. of three independent experiments, each set in triplicate.

Considering the membrane localization of the PaPlaB homolog from *L. pneumophila*, [19], we suspected a membrane localization of PaPlaB. This was confirmed by Western blot detection of PaPlaB only in the membrane fraction of *E. coli* C43(DE3) pET22-*paplaB* sedimented upon ultracentrifugation, while no PaPlaB was detected in the soluble fraction containing periplasmic and cytoplasmic proteins (Fig. 2). We next investigated whether PaPlaB is associated with inner or outer membranes of *E. coli* by separation of these two membranes using ultracentrifugation in a sucrose density gradient. Analysis of UV absorbance (A_280nm_) through the gradient after the centrifugation suggested an efficient separation of inner and outer membranes, which we assigned to be fraction 5 (inner membranes) and fractions 10-11 (outer membranes) (Fig. 3a). The refractometric measurement showed that the sucrose concentration in fractions 5 and 11 was 45 and 67 % (w/v), respectively, which is in agreement with the literature [54]. We confirmed that fractions 10-11 contain the outer membrane proteins by immunodetection of outer membrane protein TolC from *E. coli*, whereas fraction 5 did not contain *E. coli* TolC (Fig. 3b). Immunodetection of PaPlaB revealed a weak PaPlaB signal in fraction 5 and a strong signal in fractions 10-11. However, the highest esterase activity was detected for fraction 5, while the enzymatic activity of the PaPlaB-enriched fraction 11 was negligibly higher than the activity of the empty vector control (Fig. 3c).

**Fig. 3:**
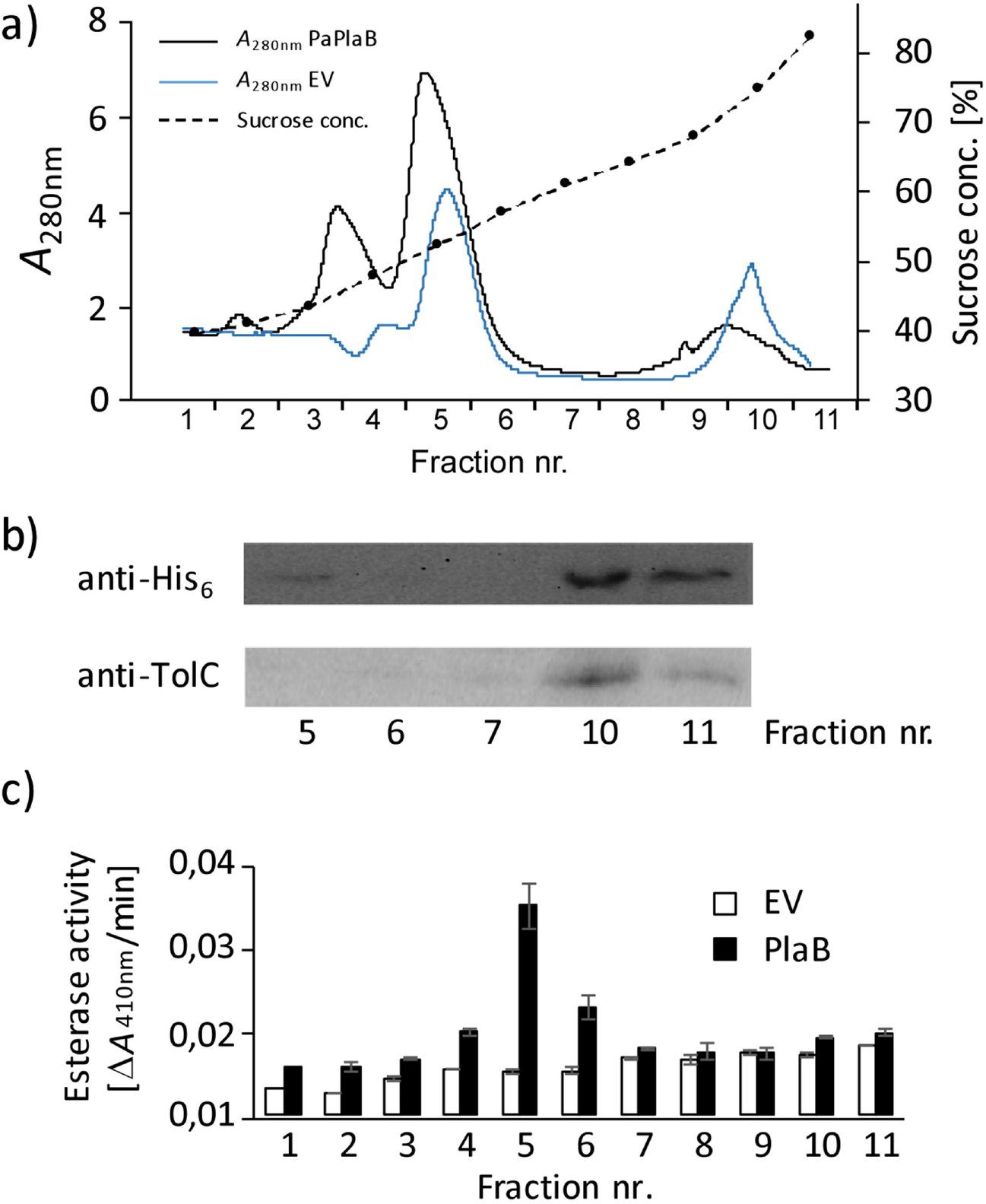
Membrane localization of PaPlaB. **a)** Isolated membranes of *E. coli* C43(DE3) pET-*paplaB* strain cultivated in LB medium (25 ml, 5 h, 37°C) were separated by sucrose density gradient. *E. coli* C43(DE3) pET22b cultivated under the same conditions was used as the empty vector (EV) control. Fractions (1 ml) were collected and their sucrose concentration was measured refractometricaly (filled circles, dashed line). Protein absorption at 280 nm is shown in solid lines. **b)** Sucrose density gradient fractions of *E. coli* C43(DE3) pET-*paplaB* were analyzed by Western blotting using the anti-His (C-term)-HRP antibody for detection of PaPlaB and primary anti-TolC antibodies combined with the anti-rabbit immunoglobulin G antibodies for detection of TolC. **c)** Enzymatic activity was measured with *p*-NPB assay by combining 10 μl of fraction and 150 μl of the substrate. The activities are means ± S.D. of two independent experiments with three samples.

For protein isolation, we used Triton X-100 detergent for extraction of PaPlaB from the membranes, while mild, non-ionic detergent DDM was added to the buffers used for IMAC purification to maintain the soluble state of PaPlaB. Elution of PaPlaB from Ni^2+^-NTA column with buffer containing an increasing concentration of imidazole resulted in highly pure PaPlaB as judged from SDS-PAGE (Fig. 4). The established protocol yielded ~0.25 mg of PaPlaB per 1 l of overexpression culture. Purified PaPlaB showed specific esterase (*p*-NPB substrate) and phospholipase A (1,2-dilauroyl phosphatidylcholine, PC_12:0_ substrate) activities of 3.41 ± 0.1 and 6.74 ± 0.8 *U*/mg, respectively.

**Fig. 4:**
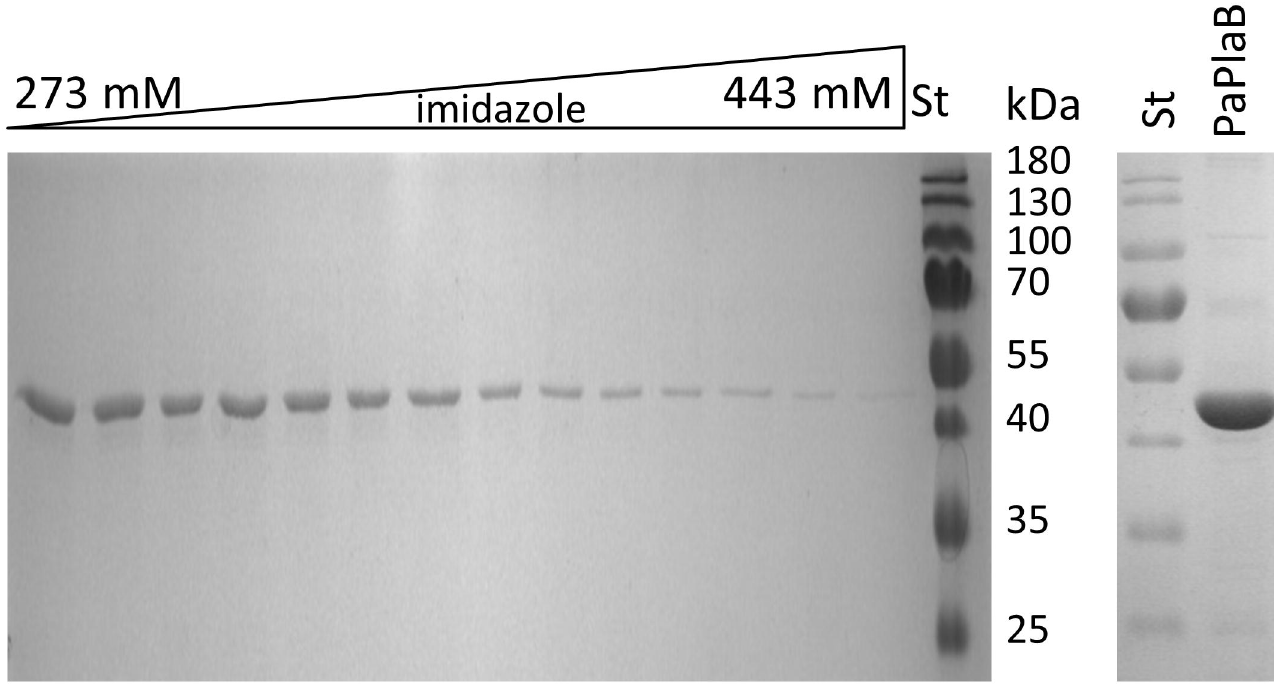
Purification of detergent-isolated PaPlaB. The fractions eluted from the Ni-NTA column (left) and pooled PaPlaB after desalting by PD-10 column (right) were analyzed by SDS-PAGE (12 % v/v). The molecular weights of protein standards (St) are indicated.

As ethylenediaminetetraacetic acid (EDTA), an inhibitor of metal-dependent enzymes, did not exert an inhibitory effect on PaPlaB (Fig. S7), we concluded that PaPlaB belongs to metal ion independent type of PLAs [55]. We furthermore examined inhibition of PaPlaB activity with two irreversible inhibitors, paraoxon and phenylmethylsulfonyl fluoride (PMSF). Under the conditions used, the activity of the paraoxon-treated PaPlaB was abolished (Fig. S7). Paraoxon covalently modifies the catalytic serine residue in the serine-hydrolase enzyme family [56]; therefore, we concluded that PaPlaB contains a nucleophilic serine in its active site, which is in agreement with the sequence-based prediction of a Ser-His-Asp catalytic triad (Fig. 1) and mutational studies (Fig. S6). Although PMSF is also utilized to identify serine hydrolases [57], it did not inhibit the PaPlaB.

Paraoxon is an organophosphate inhibitor that mimics, after the reaction with an enzyme, the first transition state. In contrast, PMSF is a sulfonyl inhibitor that mimics, after the reaction, the transition state formed by the deacylation of the acyl-enzyme in the second step of ester-hydrolysis [57]. The observed striking difference in the reactivity of paraoxon and PMSF with PaPlaB may be due to differences in the stability of the two transition state mimics.

### 3.3 PaPlaB shows promiscuous PLB and lysoPLA activities

Using an esterase activity assay, we observed that PaPlaB retained 100 % of its activity after incubation for 1 h at temperatures up to 42.5°C (Fig. S8). The thermal stability of PaPlaB was confirmed by monitoring its thermal unfolding via changes in the intrinsic fluorescence. The unfolding profile of PaPlaB revealed the transition temperature of ~53°C (Fig. S8). Hence, PaPlaB is stable and active at temperatures relevant to bacterial infections. Therefore, a temperature of 37°C was used for activity assays. We next examined the PLA activity of PaPlaB using a spectrum of glycerophospholipids (GPLs) naturally occurring in cell membranes. We showed that PaPlaB is a rather promiscuous PLA, using GPL substrates with various head groups (ethanolamine, glycerol, and choline) (Fig. 5a) and different fatty acid chain lengths (C6 – C18) (Fig. 5b). It released fatty acids from all tested substrates with specific activities in the range from 2 *U*/mg to 8 *U*/mg, with 1,2-dimyristoyl-phosphatidylethanolamine (PE_14:0_) being the best substrate. We then analyzed whether PaPlaB hydrolyses GPLs containing one fatty acid linked to the *sn*-1 position called lysoglycerophospholipids (lysoGPLs). Experiments using lysoGPLs with various head groups (ethanolamine, glycerol, and choline) showed that all three lipid types were accepted as substrates by PaPlaB (Fig. 5c). Notably, the lysoPLA activity of PaPlaB is generally lower (2 – 2.5 *U*/mg) than its PLA activity toward the respective GPLs (Fig. 5c). To analyze whether PaPlaB shows specificity for hydrolysis of fatty acids bound to *sn*-1 or *sn*-2 in GPLs, artificial fluorogenic substrates (PED-A1 for PLA1, and Red/Green BODIPY^®^ PC-A2_R/G_ for PLA2) were used. These substrates contain one alkyl group bound to glycerol by ether bond, which cannot be cleaved by PLA. Both substrates were not hydrolyzed by PaPlaB (Fig. S9), most likely due to steric hindrance caused by the bulky fluorescent dyes linked to the acyl chain of substrates. We furthermore tested whether PaPlaB hydrolyzes the natural phospholipid 1-oleoyl-2-palmitoyl-PC (PC_18:1-16:0_), which contains different fatty acids bound to glycerol. Spectrophotometric quantification of the total fatty acid amount after incubation of PaPlaB (4.3 μg/ml) with PC_18:1-16:0_ (0.5 mM) showed PaPlaB activity of 3.7 ± 0.6 *U*/mg. To identify which fatty acids were released, PaPlaB-treated PC_18:1-16:0_ samples were analyzed by GC-MS. The results of GC-MS quantification revealed 1.6 ± 0.2 μmol and 1.7 ± 0.1 μmol for palmitic and oleic acid, respectively (Table S5). This result confirmed that PaPlaB hydrolyzes both ester bonds in PC_18:1-16:0_ substrate with a similar efficiency, which classifies it into the phospholipase B (PLB) family.

**Fig. 5:**
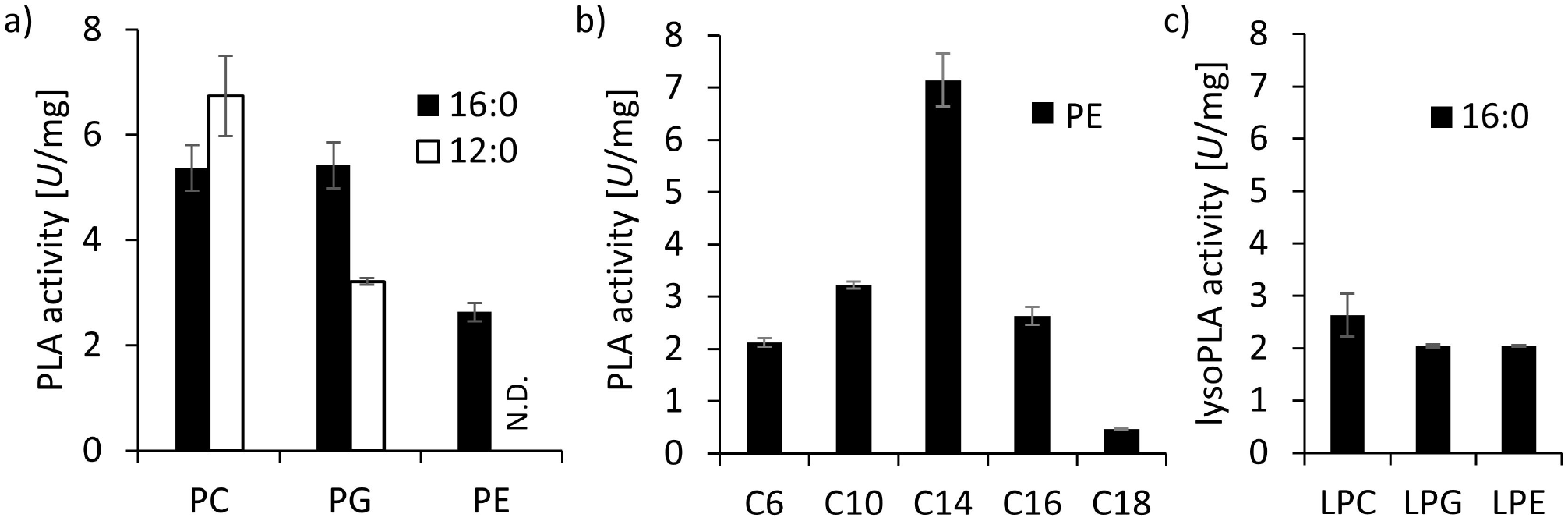
Phospholipolytic activity profile of PaPlaB. **a)** PaPlaB is a PLA that hydrolases PE, PG, and PC, which contain unsaturated FAs with C16 (16:0), and C12 (12:0) chain length commonly occurring in *P. aeruginosa* membranes. N.D. = not determined. **b)** Substrate specificity of PaPlaB measured with PE containing different FA chain lengths (C6 - C18). **c)** PaPlaB shows hydrolytic activity towards various lysophospholipids (LPE, LPG, and LPC) containing unsaturated FA with C16 (16:0) chain length. PLA and lysoPLA activities were measured by NEFA-assay using 54 ng PaPlaB per reaction. Activities are mean ± S.D. of three independent experiments with three samples.

### 3.4 PaPlaB oligomerizes in solution

Reversible formation of dimeric and tetrameric LpPlaB was observed at protein concentrations ranging from ~0.01 to 1 mg/ml. Therefore, we assessed whether purified and DDM-stabilized PaPlaB oligomerizes in solution. Size-exclusion chromatography (SEC) analysis of PaPlaB at 0.1, 0.5, and 1.0 mg/ml revealed presence of several oligomeric PaPlaB species. Protein species of ~45 kDa, and ~360 kDa, as judged through comparison with standard globular proteins of known molecular weights, were observed for all tested PaPlaB concentrations (Fig. 6a). According to the theoretical M_w_ of PaPlaB of 49.5 kDa, we interpreted these as monomeric and heptameric PaPlaB, although the exact oligomerization state cannot be reliably assessed due to detergent bound to PaPlaB and likely nonglobular shape of oligomers. Notably, at low PaPlaB concentration (0.1 mg/ml), the amount of the monomeric PaPlaB is much larger than the amount of the heptameric PaPlaB, as expected. By rising the PaPlaB concentration, the equilibrium shifts towards heptamers. Additionally, PaPlaB species of intermediate molecular masses were also detected, thus indicating a stepwise oligomerization of PaPlaB. Based on the elution volumes, we assigned these oligomers to trimeric, tetrameric, and pentameric forms of PaPlaB. At high PaPlaB concentration (1 mg/ml), pentameric PaPlaB was prevalent over tetrameric and trimeric species, while at 0.5 mg/ml predominantly tetrameric, and 0.1 mg/ml predominantly trimeric PaPlaB were observed. Thus, the equilibrium was shifted towards higher-order oligomeric species by increasing the PaPlaB concentration. The formation of homooligomers was predicted by *ab initio* docking using monomeric PaPlaB structure as an input (Fig. S10). The spatial orientation of PaPlaB monomers, dimers, and higher oligomers at the membrane suggests that PaPlaB is a peripheral membrane-bound protein (Fig. S10). This is in agreement with the negative result of the sequence-based prediction of TM helices with biological relevance (Table S6).

**Fig. 6:**
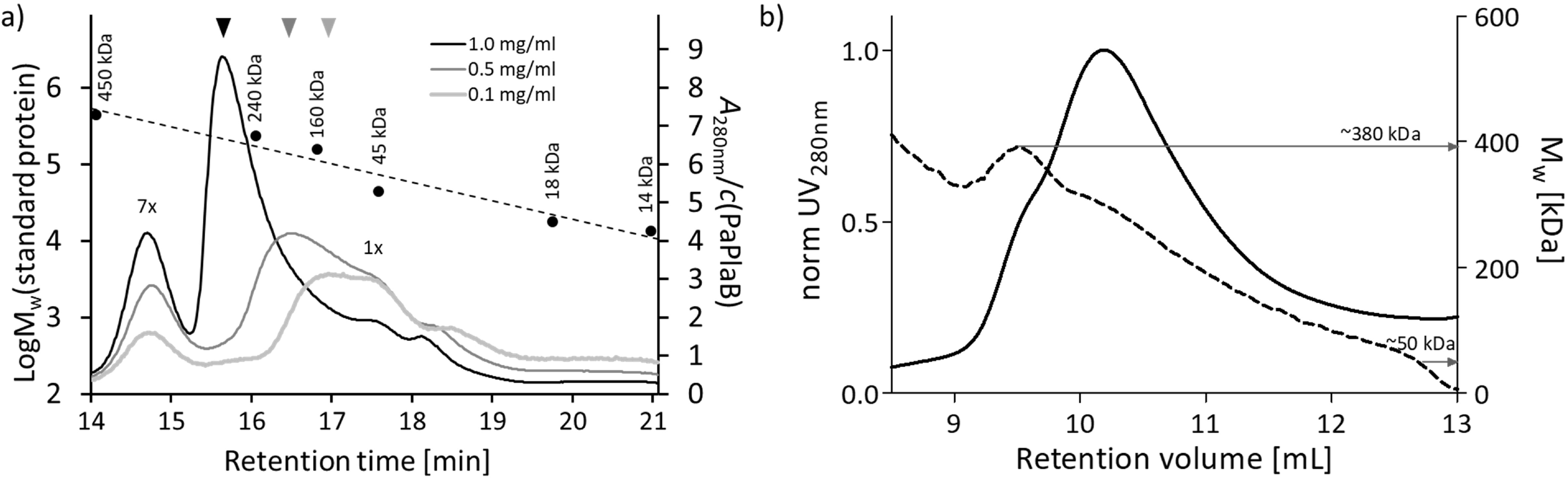
Concentration-dependent oligomerization of PaPlaB. **a)** PaPlaB (1.0, 0.5, and 0.1 mg/ml), and standard proteins (Table S3) dissolved in a buffer containing DDM were separately analyzed using Biosep-SEC-S3000 column. Proteins were detected by measuring absorbance at 280 nm (solid curves). **b)** SEC-MALS analysis using a Superdex 200 Increase column. PaPlaB (1.0 mg/ml) stabilized by DDM was detected by measuring absorbance at 280 nm (solid curve), and the overall M_w_ (dashed line) was determined with the software ASTRA 7.

Determination of absolute M_w_ of protein:detergent complexes by SEC analysis is prone to errors. Therefore we determined the absolute M_w_ of DDM-stabilized PaPlaB using multi-angle light scattering coupled to SEC (MALS-SEC). The absolute M_w_ that was determined at a concentration of 1 mg/ml revealed a distribution starting at ~380 kDa (heptamer), which was continuously decreasing to ~50 kDa (monomer) (Fig. 6b). For the PaPlaB sample with 0.1 mg/ml very broad MALS signal in the range expected for proteins with M_w_ ~50 kDa was observed (Fig. S11). Similar to SEC experiments, MALS-SEC results showed that the equilibrium of PaPlaB oligomers expectedly depends on the protein concentration.

### 3.5 PaPlaB is a major intracellular PLB of *P. aeruginosa* with hydrolytic activity towards endogenous phospholipids

To study the *in vivo* PLB function of PaPlaB in the homologous host, we constructed a *P. aeruginosa* deletion mutant *∆plaB*, which is missing the entire *plaB* gene (Fig. S1). The activity assay showed a 60 % reduction of cell-associated PLA activity in *P. aeruginosa ∆plaB* compared with *P. aeruginosa* wild-type (Fig. 7a). PLA activity of proteins secreted into the medium was not significantly different among these two strains (Fig. 7a), indicating that PaPlaB is a cell-associated and not secreted PLA of *P. aeruginosa*. PLA activity of PaPlaB demonstrated *in vitro*, and the membrane localization of the enzyme provides a hint that PaPlaB might be related to hydrolysis of cell membrane GPLs. To test this, we have isolated phospholipids (PLs) from the *P. aeruginosa* wild-type cells by extraction with an organic solvent. These PL extracts were used at 3.3 mg/ml and 0.46 mg/ml as substrates for *in vitro* PLA assay with purified PaPlaB at 450, 45, 4.5 and 0.45 ng/ml.

**Fig. 7:**
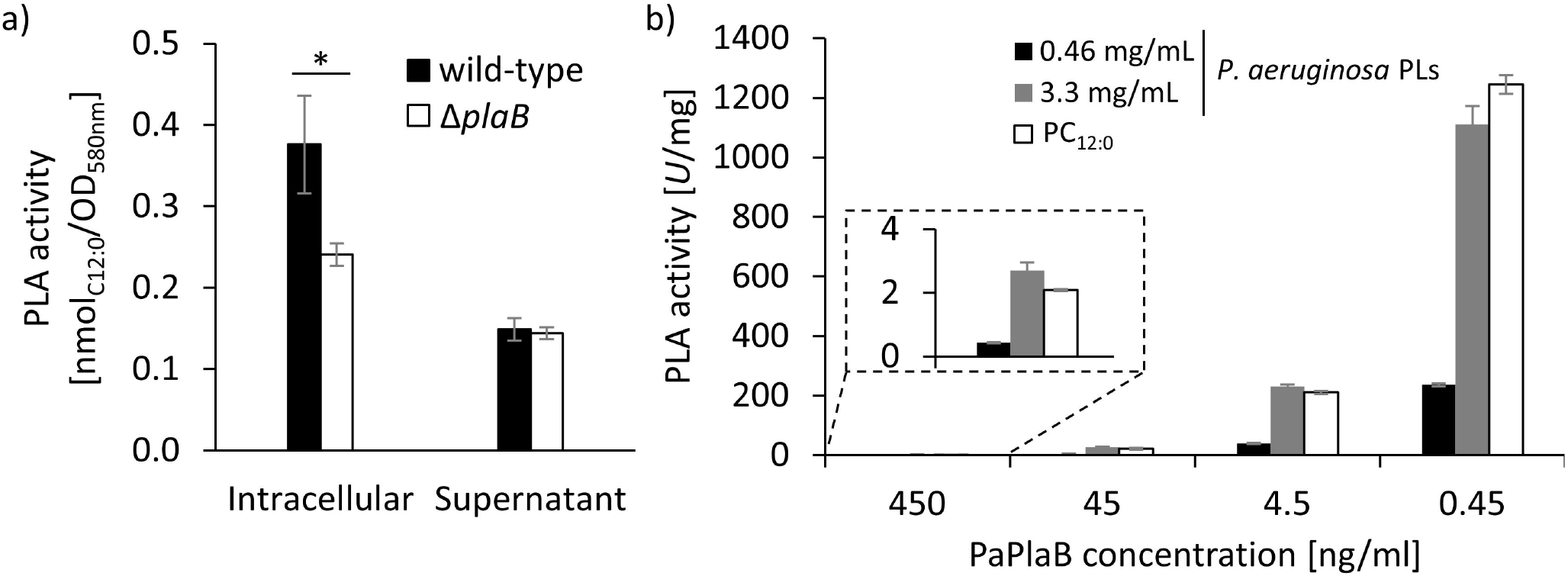
PaPlaB is an intracellular PLA of *P. aeruginosa* that releases fatty acids from endogenous phospholipids. **a)** PLA activity of the whole cells and the supernatant of *P. aeruginosa* wild-type and *∆plaB* cultivated in LB medium overnight at 37°C. Cells washed with fresh LB medium were disrupted by ultrasonication before measurement. NEFA assay with PC_12:0_ substrate (25 μl) was performed using cell lysates (25 μl) adjusted to OD_580nm_ = 10 or undiluted cell-free supernatant (25 μl). Results are the means ± S.D. of three measurements with three biological replicates. Statistical analysis was performed using the *t*-test, * p < 0.05. **b)** PLA activity of purified PaPlaB was measured by NEFA assay using endogenous PLs isolated from *P. aeruginosa* wild-type cells and synthetic PC_12:0_, which was used as control. Free fatty acids were quantified after 15 min incubation of PaPlaB with the substrate at 37°C. Activities are mean ± S.D. of three measurements with three biological replicates.

Results showed that PaPlaB hydrolyzes endogenous PLs with high efficiency (Fig. 7b). Hence, assays with 3.3 mg/ml endogenous PLs showed comparable activities to these measured with PC_12:0_, which was among the best PaPlaB substrates. PaPlaB activity with endogenous PLs was higher at higher substrate concentrations as expected for enzyme-catalyzed reactions. We furthermore observed that specific PaPlaB activities immensely increase by diluting the PaPlaB samples. Consequently, 2 and > 1100 *U*/mg activities were respectively measured with 450 and 0.45 ng/ml enzyme and 3.3 mg/ml endogenous PLs. We confirmed the activation of PaPlaB by dilution in the assays performed with 0.46 mg/ml endogenous PLs or PC_12:0_ (Fig 7b).

### 3.6 PaPlaB affects biofilm amount and architecture

To investigate whether PaPlaB affects the formation, maturation, and dispersion of biofilm, we have performed long-time studies (8 – 216 h) of biofilm formation in microtiter plates (MTP) under static conditions (crystal violet assay) and in the chamber with a continuous supply of the nutrients under dynamic conditions (confocal laser scanning microscopic (CLSM) analysis) (Figs. 8a and 8b). *P. aeruginosa ∆plaB* produces significantly less biofilm under static conditions than the wild-type strain after 8, 24, 48, and 72 h of growth, indicating that PaPlaB plays a role in initial attachment and maturation [58, 59] of *P. aeruginosa* biofilm (Fig. 8a). Under these conditions, the biofilm amount in *P. aeruginosa ∆plaB* and wild-type cultures grown for 6 and 9 days showed no significant difference, indicating that PaPlaB likely does not have a function for biofilm dispersion. Based on these results, we examined the biofilm assembly of 24, 72, and 144 h-old biofilms by using CLSM [60]. Large differences between the *P. aeruginosa ∆plaB* and WT were observed (Figs. 8b and S11). After 72 h, the wild-type strain forms larger aggregates opposite to small-sized aggregates observed for the *∆plaB* strain. The lower density of 72 h-old biofilms found by CLSM correlates with less biofilm quantified by crystal violet assay after 72 h of growth. Interestingly, although the crystal violet assay did not reveal significant differences after six days of growth, the CLSM showed differences. Hence, the wild-type nearly homogeneously and densely covered the surface of the flow-cell coverslip after 144 h, whereas the *∆plaB* strain showed less dense coverage indicating impaired maturation [58, 59].

**Fig. 8:**
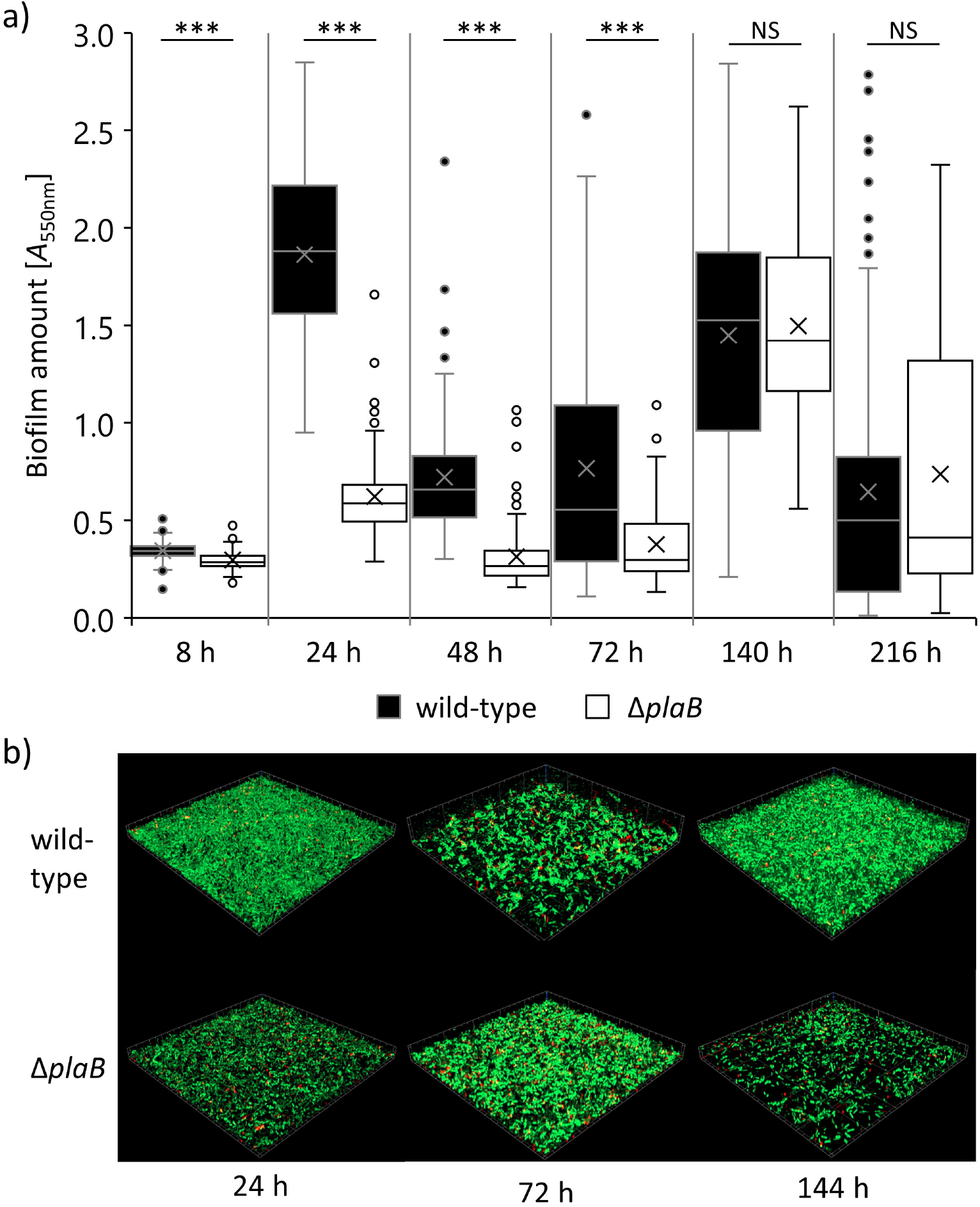
PaPlaB affects biofilm formation in *P. aeruginosa*. **a)** *P. aeruginosa* wild-type and *∆plaB* were cultivated in 96-well MTP (LB medium, 37°C, without aeration). The cells not attached to the plastic surface were removed, and the biofilm stained with crystal violet was quantified at 550 nm. The results are mean ± S.D. of three independent experiments with five biological replicates each measured eight times. Statistical analysis was performed using the *t*-test, *** p < 0.001. **b)** Biofilm architecture analyzed by CLSM after 24, 72, and 144 h growth at 37°C in a flow cell with continuous supply (50 μl/min) of LB medium. Experiments were repeated two times, each with one biological replicate that was analyzed at three different points by imaging a section of 100 x 100 μm. All collected images are shown in Fig. S12.

## 4. Discussion

Here we identified *P. aeruginosa* PA01 gene *pa2927*, which encodes a novel PLB (PaPlaB) with a function in biofilm assembly. This enzyme shows moderate global sequence homology (Fig. 1a) with a known virulence-related outer membrane PLA (LpPlaB) of *L. pneumophila* [18, 19, 46, 47] Sequence alignment of LpPlaB and PaPlaB revealed strongly conserved catalytic triad residues (Ser, His, Asp) embraced in the N-terminal domain which is predicted to resemble α/β-hydrolase fold (Fig 1b). By sequence alignment predicted and by mutational analysis confirmed (Fig. S6) catalytic triad resembles in the three-dimensional model the catalytic triad reported in other Ser-hydrolases with PLA activity [53]. C-terminal sequences, in contrast to N-terminal, are not similar within PlaB-family and do not show similarity to other proteins (Fig. 1a). It is thought that the C-terminus of the PlaB-family proteins folds into a distinct domain [18]. This was proposed for LpPlaB [18] and is suggested from *de novo* structural model of PaPlaB (Fig. 1b). The functional importance of the C-terminal domain was suggested, as 15 C-terminal residues were shown to modulate phospholipolytic A activity of LpPlaB [46], although the reason for this is unknown. Interestingly, PaPlaB is missing these 15 C-terminal residues (Fig. S2) what might be related to subtle catalytic differences between LpPlaB and PaPlaB. For example, PaPlaB has comparable PLA activities with PG and PC substrates (Fig. 5a), while LpPlaB hydrolyses PG two times faster than PC [18, 19].

We next analyzed whether PaPlaB is an intracellular or extracellular protein because the physiological function of secreted bacterial PLAs and PLBs differ substantially from the function of intracellular enzymes [7, 61]. Extracellular PLA/Bs are toxins involved in host cell membrane disruption [62] or modulation of host cell pathways through the release of bioactive compounds [7]. On the other hand, the function of intracellular PLA/Bs in bacteria is still not clearly established although we recently discovered novel cytoplasmic membrane-bound PLA1 PlaF from *P. aeruginosa* which remodeling of membrane GPLs is suggested as a virulence mechanism [9, 10]. Interestingly, the function of intracellular PLAs for the regulation of fatty acyl chain composition in GPLs through a deacylation-reacylation pathway called Lands' cycle was described in yeast [63] and other eukaryotes [64].

Using *P. aeruginosa ∆plaB* we observed a ~60 % reduction of a cell-associated PLA activity compared to the wild type, whereas extracellular PLA activities did not significantly differ (Fig. 7a). These results suggest that PaPlaB is the main intracellular PLA of *P. aeruginosa*. In agreement with this result is observed membrane localization of catalytically active PaPlaB recombinantly produced in *E. coli* C43(DE3) [32] (Fig. 2), although a large portion was accumulated in catalytically inactive aggregates. Furthermore, activity assays and Western blot analysis of sucrose density gradient fractionated membranes isolated from fragmented *E. coli* C43(DE3) cells overexpressing PaPlaB indicated dual membrane localization of PaPlaB (Fig. 3). Yet, only the cytoplasmic membrane fraction showed PaPlaB activity while the activity of the outer membrane fraction was comparable to the empty vector control strain. These results are only partially agreeing with the suggested outer membrane localization of LpPlaB, [46] because LpPlaB showed the highest PLA activity in outer membrane Momp protein-enriched fractions of *L. pneumophila*. However, it also showed substantial activity (~70 % of outer membrane activity) in the fractions containing inner membranes [46]. The drawback of this fractionation method is the difficulty of absolute separation of outer from inner membranes, which was described by several research groups [46, 65, 66]. Keeping in mind that LpPlaB and PaPlaB do not have predicted TM helix or β-barrel-like structure, which were recognized in all hitherto known integral membrane proteins [67, 68], it is likely that these hydrophobic proteins are peripherally associated with one or both membranes. In line with this suggestion is our observation that monomeric PaPlaB in the presence of DDM micelles (~70 kDa large [69]) has the M_w_ of ~45 kDa (estimated by SEC, (Fig. 6a)) which is close to the theoretical M_w_ (49.5 kDa). Hence, it seems that PaPlaB has not been embedded into DDM micelle, as it would be expected for a true integral membrane protein. Additionally, the prediction of the orientation of PaPlaB at the membrane using the structural model suggests its peripheral interaction with the membrane (Fig. S10).

These findings strengthen our hypothesis that PaPlaB is rather a peripheral membrane protein. Interestingly, in PaPlaB and LpPlaB, [46] no recognizable signature for their secretion across the membrane was found; therefore, it remains unknown how these proteins are targeted across and to the membrane. Further experiments are needed to clarify cellular localization and the membrane anchoring mechanism of LpPlaB and PaPlaB.

The additional similarity of LpPlaB and PaPlaB is the phenomenon that these proteins homooligomerize at high concentrations, which is accompanied by a decrease in their enzyme activity (Fig. 7b) [47]. Using the SEC method, we observed the equilibrium of PaPlaB monomers and several oligomeric species with M_w_ of up to ~360 kDa, likely heptamers, at concentrations 0.1, 0.5, and 1.0 mg/ml. Keeping in mind that the shape, which presumably deviates from a sphere (Figs. 1b and S10), and bound DDM molecules make a difficult precise determination of oligomeric state by SEC, we have determined an absolute M_w_ of PaPlaB by MALS method. MALS analyses confirmed PaPlaB monomers and the formation of various oligomers of up to ~380 kDa (Fig. 6b). The observation that PaPlaB:DDM species of M_w_ between ~45 and ~380 kDa were simultaneously present in the same sample suggests a stepwise oligomerization of PaPlaB. This is in agreement with the structural prediction of PaPlaB oligomers containing from 3 to 7 PaPlaB monomers.

SEC and MALS results revealed that increasing the PaPlaB concentration leads to the enrichment of higher oligomeric species. This is similar to LpPlaB, for which were identified only homotetramers at the concentration of ≥ 0.3 mg/ml and the mixture of tetramers and dimers at protein concentrations ≤ 0.05 mg/ml by analytical ultracentrifugation [47]. Furthermore, the oligomerization at higher protein concentrations was accompanied by a strong decrease in activities of PaPlaB and LpPlaB several hundredfolds (Fig. 7b) [47], which was suggested as a mechanism of protecting the host from uncontrolled degradation of own membranes [47]. The activity of *P. aeruginosa* phospholipase A ExoU [70], and human PLA_2_ [71], has been suggested to be regulated through homomeric protein:protein interactions. However, further studies are necessary to elucidate the molecular mechanism of PaPlaB activation and particularly the role of the membrane for it.

Although LpPlaB and PaPlaB seem not to be essential for bacterial life (Fig. S1), they both affect the important virulence properties of their hosts. It was suggested that the regulation of intracellular replication of *L. pneumophila* is a mechanism of LpPlaB-mediated virulence [46], while the regulation of biofilm maturation was suggested as a mechanism of PaPlaB-mediated virulence (Fig. 8). Despite the exact molecular mechanism by which LpPlaB and PaPlaB contribute to bacterial virulence is unknown, it is most likely that phospholipid-degrading activities are essential for their virulence function. We have shown that PaPlaB rapidly hydrolyses PE (Fig. 5b), which is the most abundant bacterial GPL [72], at the same rate as it hydrolysed GPLs extracted from the membranes of *P. aeruginosa* (Fig 7b). We could show that the biochemical function of PaPlaB is related to the complete deacylation of GPLs to fatty acids and glycerophosphoalcohol as shown by lysoPLA assay (Fig. 5c) and GC-MS analysis of fatty acid products released from PC_18:1-16:0_ (Table S5).

In conclusion, the ability of PaPlaB to rapidly degrade endogenous GPLs and above-proposed membrane localization of this novel PLB of *P. aeruginosa*, suggesting that PaPlaB might modulate the molecular GPL profile of *P. aeruginosa* membranes similarly as described for PlaF [9, 10], PLA_2_ from rat [73], yeast [63] and other eukaryotes [64]. This adaptive GPL modulation might indirectly affect biofilm formation of *P. aeruginosa* what goes along with findings that *P. aeruginosa* undergoes drastic changes in membrane GPL composition upon transition from the planktonic to a biofilm lifestyle [72]. Our results contribute to a still limited understanding of the virulence mechanism of PLA/B from pathogenic bacteria that may represent a previously not explored family of antibiotic targets.

## Acknowledgment

This study was funded by the Deutsche Forschungsgemeinschaft (DFG, German Research Foundation) – project number 267205415 – CRC 1208 (project A01 to LS, A02 to FK, A03 to HG and A10 to AK). HG is grateful for computational support by the “Zentrum für Informations und Medientechnologie” at the Heinrich-Heine-Universität Düsseldorf and the computing time provided by the John von Neumann Institute for Computing (NIC) to HG on the supercomputer JUWELS at Jülich Supercomputing Centre (JSC) (user IDs: HKF7; HDD18). We thank Christoph Strunk and Esther Knieps-Grünhagen (Heinrich Heine University Düsseldorf, IMET) for their help with the generation of the expression plasmid and SEC analysis, respectively, Muttalip Caliskan (Heinrich Heine University Düsseldorf, IMET) for providing GPLs extract, and Prof. Karl-Erich Jaeger (Heinrich Heine University Düsseldorf, IMET) for valuable discussions.

