## Supplemental information for "Novel intracellular phospholipase B from *Pseudomonas aeruginosa* with activity towards endogenous phospholipids affects biofilm assembly"

**Table S1:** Oligonucleotides used in this study.

| **Name** | **DNA Sequence (5ʹ🡪3ʹ)** |
| --- | --- |
| paplaB_fw | GGAATTCATATGCCCCGTTCGATCGTCATCG |
| paplaB_rv | GGAATTGAGCTCTCA**GTGGTGGTGGTGGTGGTG**GGGGTTCCTGAAGACGAATACG |
| *plaB*_S79A_fw | CGGTTGACGTGGTGGTGCACgccACCGGCACCCTGGTGCTGCG |
| *plaB*_D196A_fw | AGGCCGGCAGCGcCGGCACTGTGCGGATCGCCACGGCGAACC |
| *plaB*_H244A_fw | CATCGTCGACGGCGAGAACgcCGCCAGCGTGGCGCTGAAGG |
| *plaB*_S79A_rv | CGCAGCACCAGGGTGCCGGTGGCGTGCACCACCACGTCAACCG |
| *plaB*_D196A_rv | GGTTCGCCGTGGCGATCCGCACAGTGCCGGCGCTGCCGGCCT |
| *plaB*_H244A_rv | CCTTCAGCGCCACGCTGGCGgcGTTCTCGCCGTCGACGATG |

*Nde*I and *Sac*I restriction sites are underlined and the sequence encoding a hexahistidine-tag is indicated bold.

**Table S2:** Glycerophospholipid and lysoglycerophospholipid substrates used in activity assays.

| Name | Abbreviation |
| --- | --- |
| 1,2-dipalmitoyl-sn-glycero-3-phosphocholine | PC_16:0_ |
| 1,2-dilauryl-sn-glycero-3-phosphocholine | PC_12:0_ |
| 1,2-dipalmitoyl-sn-glycero-3-phospho-(1'-rac-glycerol) | PG_16:0_ |
| 1,2-dilauryl-sn-glycero-3-phospho-(1'-rac-glycerol) | PG_12:0_ |
| 1,2-dicaproyl-sn-glycero-3-phosphoethanolamine | PE_6:0_ |
| 1,2-dicapryl-sn-glycero-3-phosphoethanolamine | PE_10:0_ |
| 1,2-dilauyl-sn-glycero-3-phosphoethanolamine | PE_12:0_ |
| 1,2-dipalmitoyl-sn-glycero-3-phosphoethanolamine | PE_16:0_ |
| 1,2-distearyl-sn-glycero-3-phosphoethanolamine | PE_18:0_ |
| 1‑palmitoyl-2-hydroxy-sn-glycero-3-phosphocholine | LPC_16:0_ |
| 1-palmitoyl-2-hydroxy-sn-glycero-3-phospho-(1'-rac-glycerol) | LPG_16:0_ |
| 1-palmitoyl-2-hydroxy-sn-glycero-3‑phosphoethanolamine | LPE_16:0_ |
| 1-oleoyl-2-palmitoyl-sn-glycero--3-phosphocholine | PC_18:1-16:0_ |

**Table S3:** Standard proteins used for size-exclusion chromatographic analysis.

| Standard protein | Retention time [min] | Molecular weight [Da] |
| --- | --- | --- |
| Ribonuclease A | 20.714 | 13,700 |
| Myoglobin | 19.597 | 17,800 |
| Albumin egg | 17.642 | 45,000 |
| Aldolase rabbit | 16.947 | 160,000 |
| Catalase bovine | 16.259 | 240,000 |
| Ferritin horse | 14.444 | 450,000 |
| Uridine | 22.955 | 244 |

**Table S4:** CLSM settings.

|  | Propidium iodide | SYTO 9 |
| --- | --- | --- |
| Gamma value | 1.0 | |
| Pinhole | 51 µm | |
| Detector digital | 1.0 | |
| Laser intensity | 0.2 % | |
| Laser wavelength | 561 nm | 488 nm |
| Detection wavelength | 460-700 nm | 410-560 nm |
| Detector gain | 650 V | 600 V |

**Table S5:** GC-MS quantification of oleic (C18:1) and palmitic (C16:0) fatty acid released by PaPlaB from PC 18:1-16:0 substrate.

| Sample | Measurement* | n(C_16:0_) [µmol] | n(C_18:1_) [µmol] |
| --- | --- | --- | --- |
| 1 | 1 | 1.32 | 1.57 |
| 1 | 2 | 1.38 | 1.62 |
| 2 | 1 | 1.69 | 1.78 |
| 2 | 2 | 1.83 | 1.89 |
| 3 | 1 | 1.59 | 1.54 |
| 3 | 2 | 1.61 | 1.55 |
| Average | Average | 1.57 | 1.66 |
| S.D. | S.D. | 0.19 | 0.14 |

*Each of three PaPlaB PC 18:1-16:0 extracts was measured two times.

**Table S6:** Prediction of TM helices in PaPlaB using TMpred server

| Starting residue | Ending residue | Score* |
| --- | --- | --- |
| 168 | 186 | 710 |
| 367 | 386 | 434 |
| 165 | 183 | 384 |
| 357 | 376 | 2 |

*Only scores above 500 are considered significant

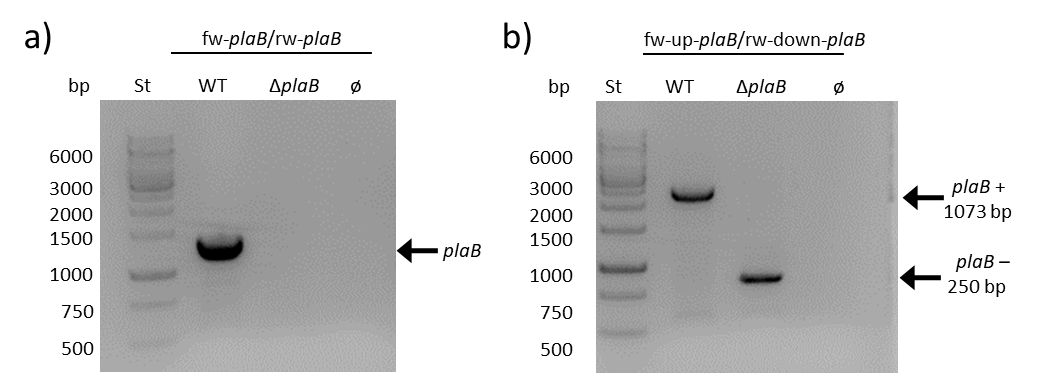

**Fig. S1: PCR verification of *P. aeruginosa* Δ*plaB* strain.** a) Agarose gel electrophoretic analysis of PCR amplification products using fw-*plaB* (5´-GGAATTGAGCTCTCAGGGGTTCCTGAAGACGAATAC-3´) and rw-*plaB* (5´-CTCTAGAGATGCCCCGTTCGATCGTCATC-3´) oligonucleotides that bind the first and last 20 bp of *paplaB* yielding the DNA product of 1332 bp for the *P. aeruginosa* wild-type were used. For the *P. aeruginosa* Δ*plaB* strain DNA product is not expected. b) Agarose gel electrophoretic analysis of PCR amplification products using fw-up-*plaB* (5´-AGAAGGTCGAACGGCAGTATC-3´) and rw-down-*plaB* (5´-GCGTTGTACCGTCGCTATG-3´) oligonucleotides that bind 668 bp upstream and 415 bp downstream of *paplaB* gene, respectively*.* DNA products of 2415 bp (*paplaB* + 1073 bp) and 1082 bp (*paplaB* - 250 bp) for the *P. aeruginosa* wild-type and Δ*plaB* strains are respectively expected*.* Ø represents the control reaction in which water was used instead of the DNA template.

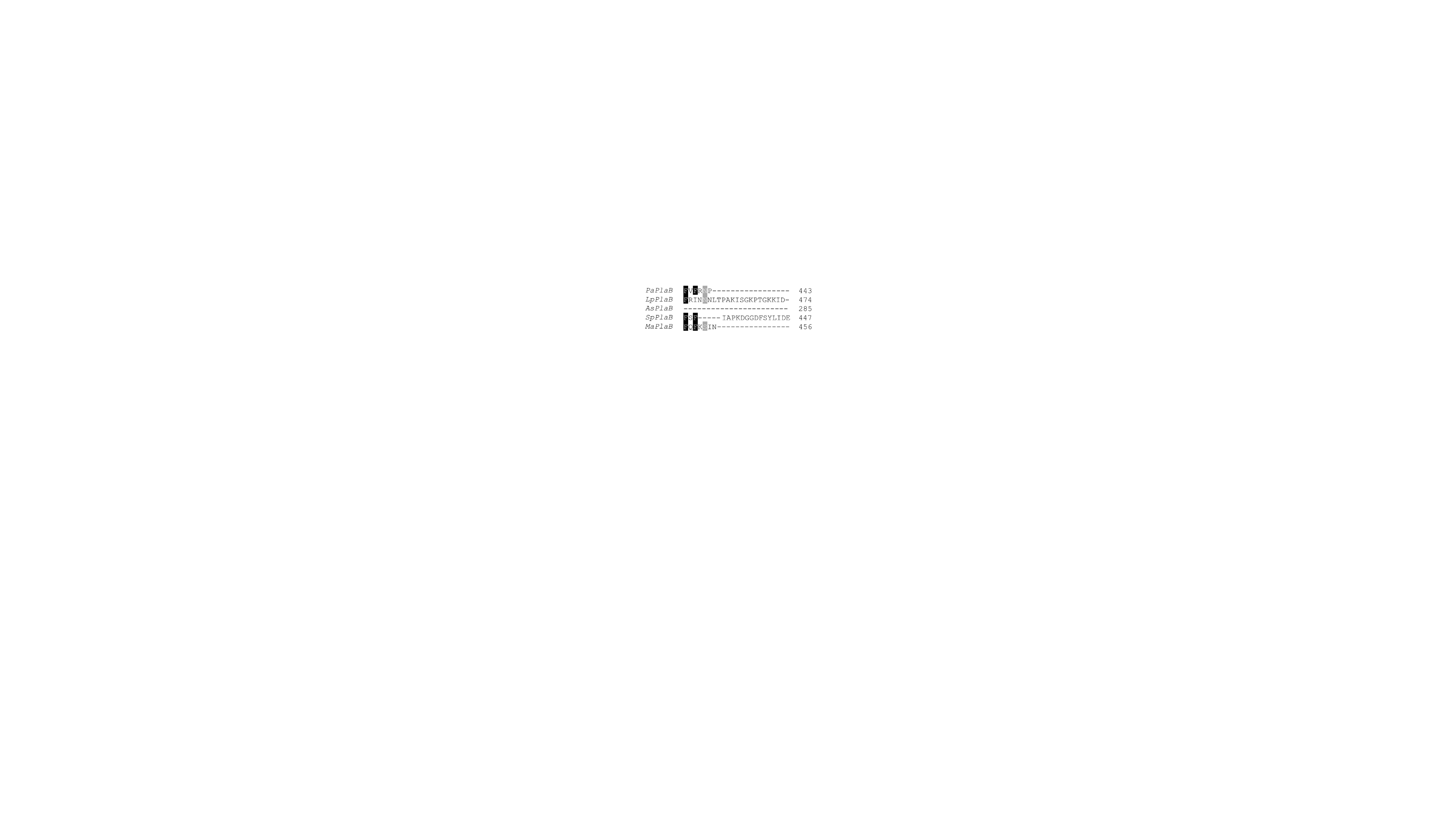

Fig. S2: Sequence alignment of PaPlaB from *P. aeruginosa* (PaPlaB), *Legionella pneumophila* (LpPlaB), *Alishewanella sp.32-51-5* (AsPlaB), *Shewanella pealeana* (SpPlaB), and *Marinobacter algicola* (MaPlaB). Here are shown C-terminal residues, which are missing in figure 1a. Black and grey background indicate identical and similar residues in at least three proteins, respectively.

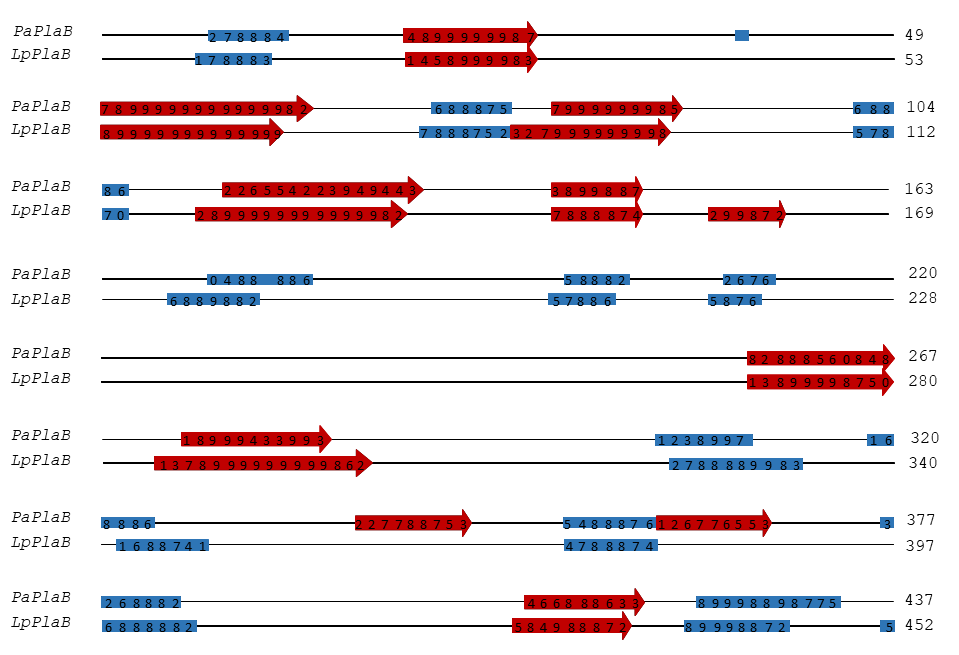

Fig. S3: Predicted secondary structure elements for PaPlaB and LpPlaB. Numbers in red arrows (α-helices) and blue boxes (β-strands) represent the JPred4^1^ secondary structure prediction score (1 = unlikely, 10 = highly probable) for each residue.

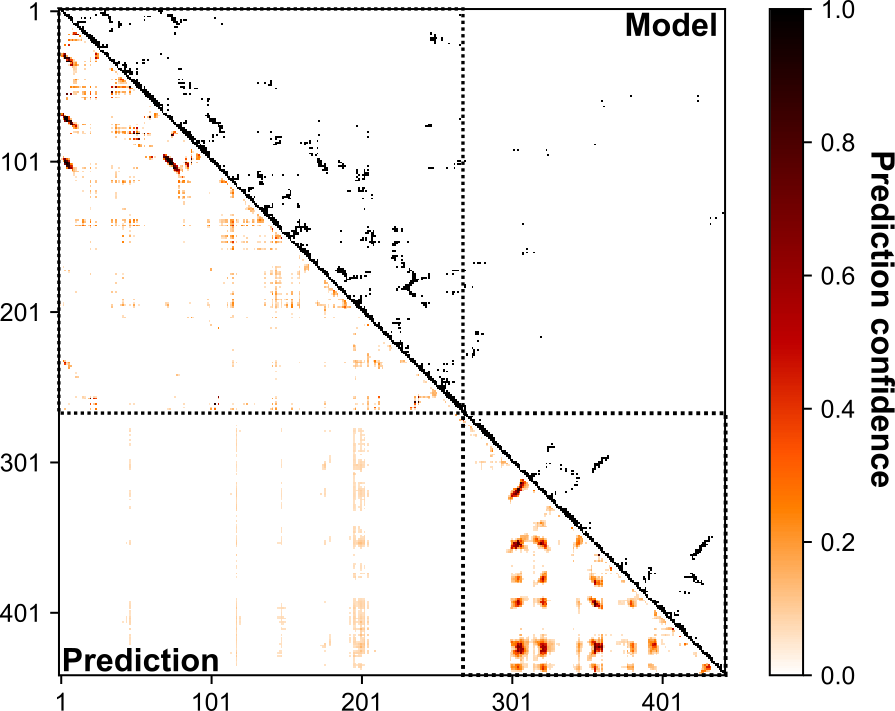

**Fig. S4: Prediction of the two-domain architecture of PaPlaB.** Contact map obtained from MetaPSICOV2 residue-residue contact predictions (lower triangle; colored according to the prediction confidence) and from the Phyre2 structural model (upper triangle; black dots). The predicted contact map shows two contact clusters (marked by dashed squares) located in the N- and C-termini, respectively, suggesting a two-domain structure.

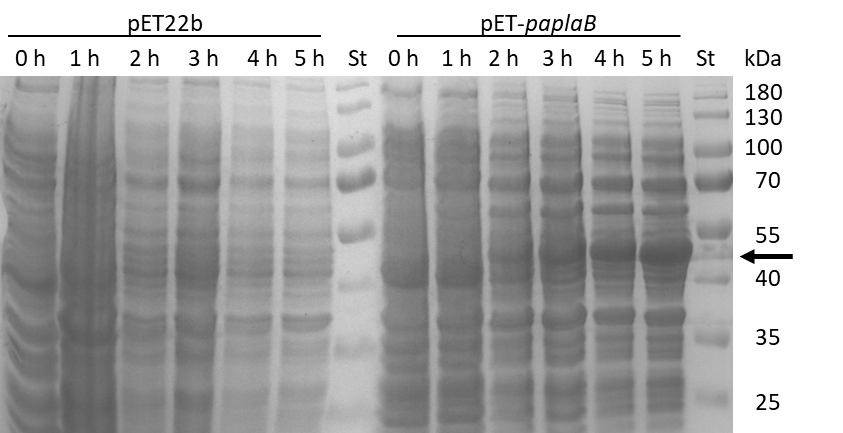

**Fig. S5: SDS-PAGE of heterologously expressed PaPlaB.** Overexpression of PaPlaB in *E. coli* C43(DE3) carrying pET-*paplaB* was induced by IPTG, and samples (10 µl, OD_580nm_ = 10) collected before expression (0 h) and 1, 2, 3, 4, and 5 h after induction were analyzed by SDS-PAGE (12 % v/v). Molecular weights of protein standards (St) are indicated (right). PaPlaB is indicated with the black arrow.

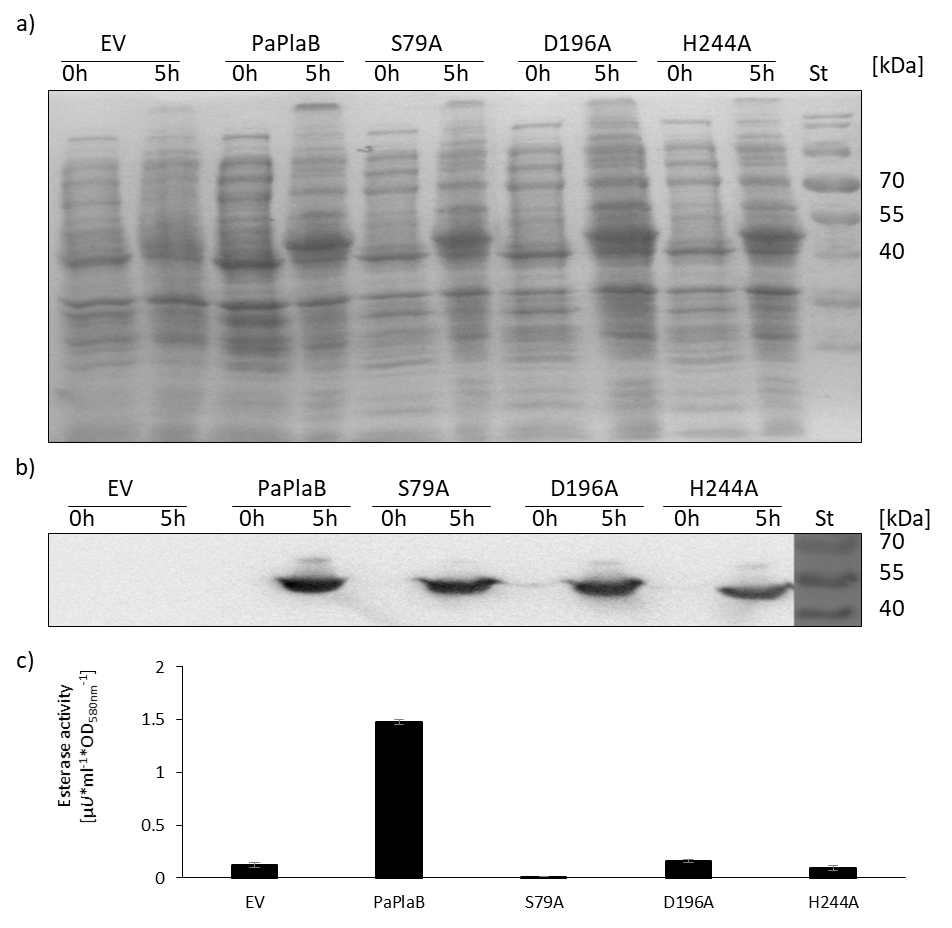

Figure S6: Mutation of PaPlaB active site residues. a) Coomassie stained SDS-PAGE (12% (w/v)) analysis of expression of *paplaB* variants in *E. coli* C43(DE3) carrying the expression plasmids pET22b-*paplaB*_S79A_, pET22b-*paplaB*_D196A_, pET22b-*paplaB*_H244A_ was induced with 1 mM IPTG followed by cultivation at 37 °C. c) Immunological detection of PaPlaB variants by Western blot using the anti-His(C-term)-HRP conjugate antibodies. *E. coli* C43(DE3)-pET22b-*paplaB*_h6_ was tested as positive control and *E. coli* C43(DE3)-pET22b as negative control under the same conditions. For analyses in b) and c) was used 10 µl of the cell lysate (O.D._580nm_ of 20). Molecular weights of standard proteins (St) are shown on the right. c) Esterase activity of cell lysates collected before induction and 5h after induction was measured using pNPB substrate. Results are the mean values and standard deviations of three measurements with the same biological replicate.

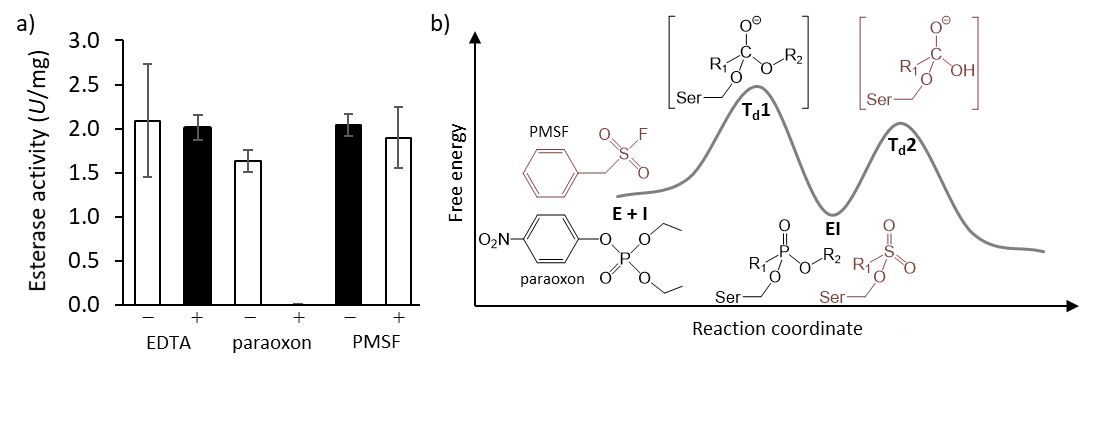

Fig S7: Inhibition of PaPlaB. Residual esterase activities were determined after incubation of PaPlaB (1.21 µM) with 10 mM EDTA, 1 mM paraoxon or 1 mM PMSF at 30ºC for 1.5 h. Inhibited PaPlaB samples (2.5 µl) were used for the determination of esterase activity with the *p*-NPB substrate (100 µl) at 37°C. Non-inhibited control samples (- inhibitor) contained PaPlaB treated with the propane-2-ol (for PMSF and paraoxon) or 100 mM Tris-HCl, pH 8 (for EDTA). The results are mean ± S.D. of two independent experiments with three samples.

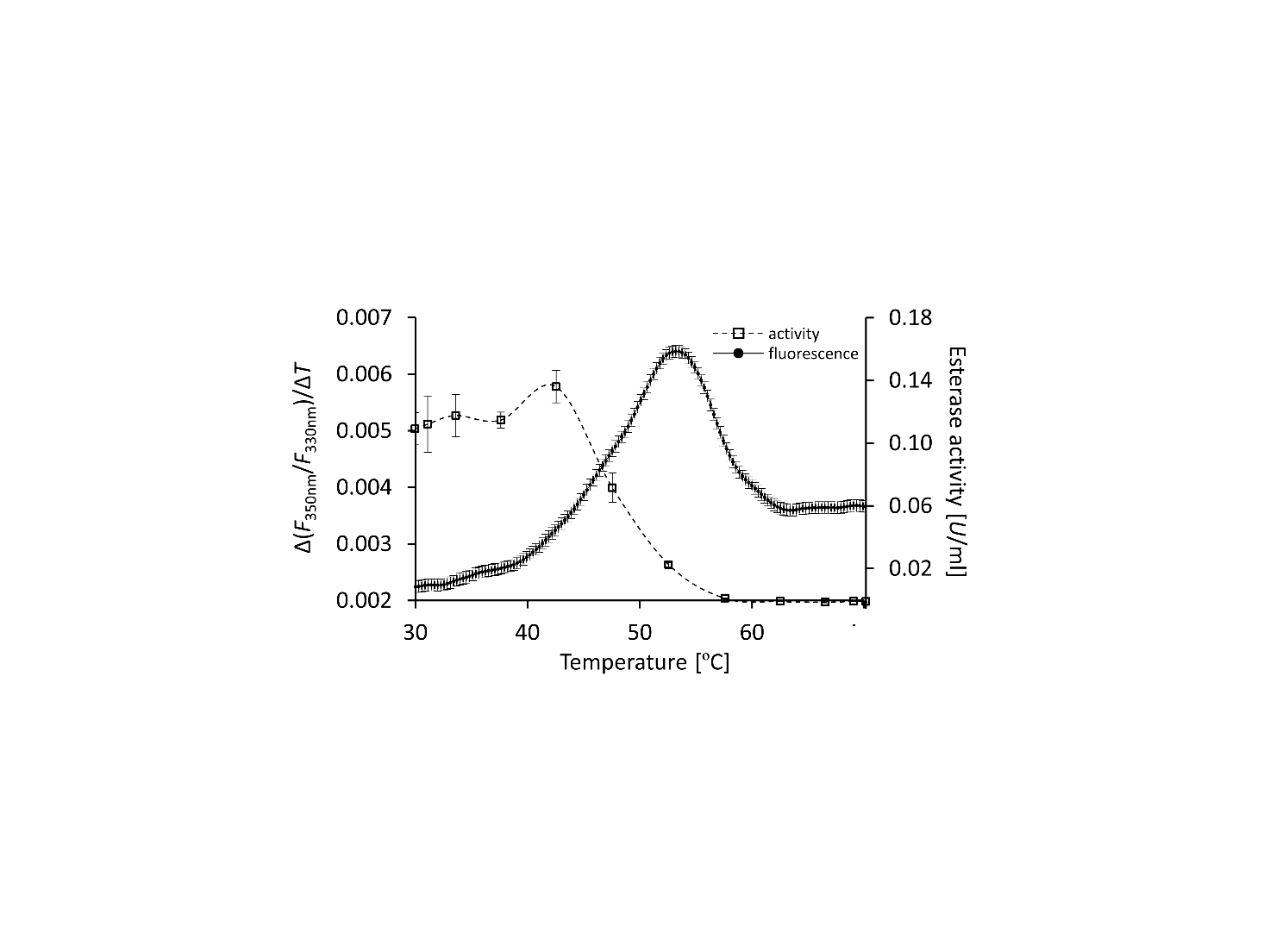

Fig. S8: Thermal stability of the detergent-stabilized PaPlaB. Thermal inactivation and denaturation of PaPlaB suggest its high thermal stability. Dashed line: Temperature-dependent PaPlaB stability determined enzymatically after incubating PaPlaB (0.24 mg/ml) at different temperatures (30-70°C) for 1 h followed by measurement of esterase activity with 2.4 mg PaPlaB using the *p*-NPB assay. Solid line: Change in intrinsic tryptophan fluorescence of PaPlaB measured by differential scanning fluorometry (nanoDSF). Esterase activities and nanoDSF results are mean ± S.D. of three measurements.

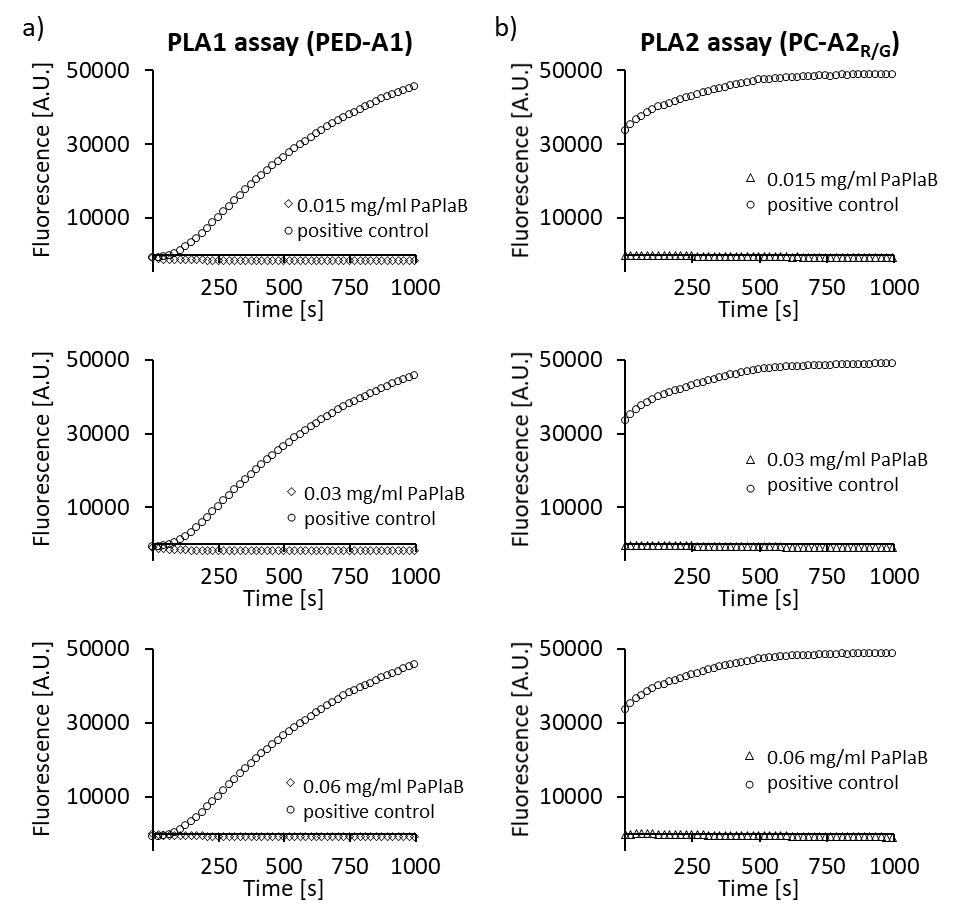

Fig. S9: PLA1 (a) and PLA2 (b) activities of PaPlaB were measured fluorimetrically using artificial PED-A1 and PC-A_R/G_ substrates containing acyl chains labeled with a fluorescent dye. Substrate and PaPlaB at three different concentrations were combined in 96-well MTP, and fluorescence was measured at room temperature. The control enzymes were PLA1 of *Thermomyces lanuginosus* and PLA2 of *Naja mocambique*. Activities are means ± S. D. of three measurements.

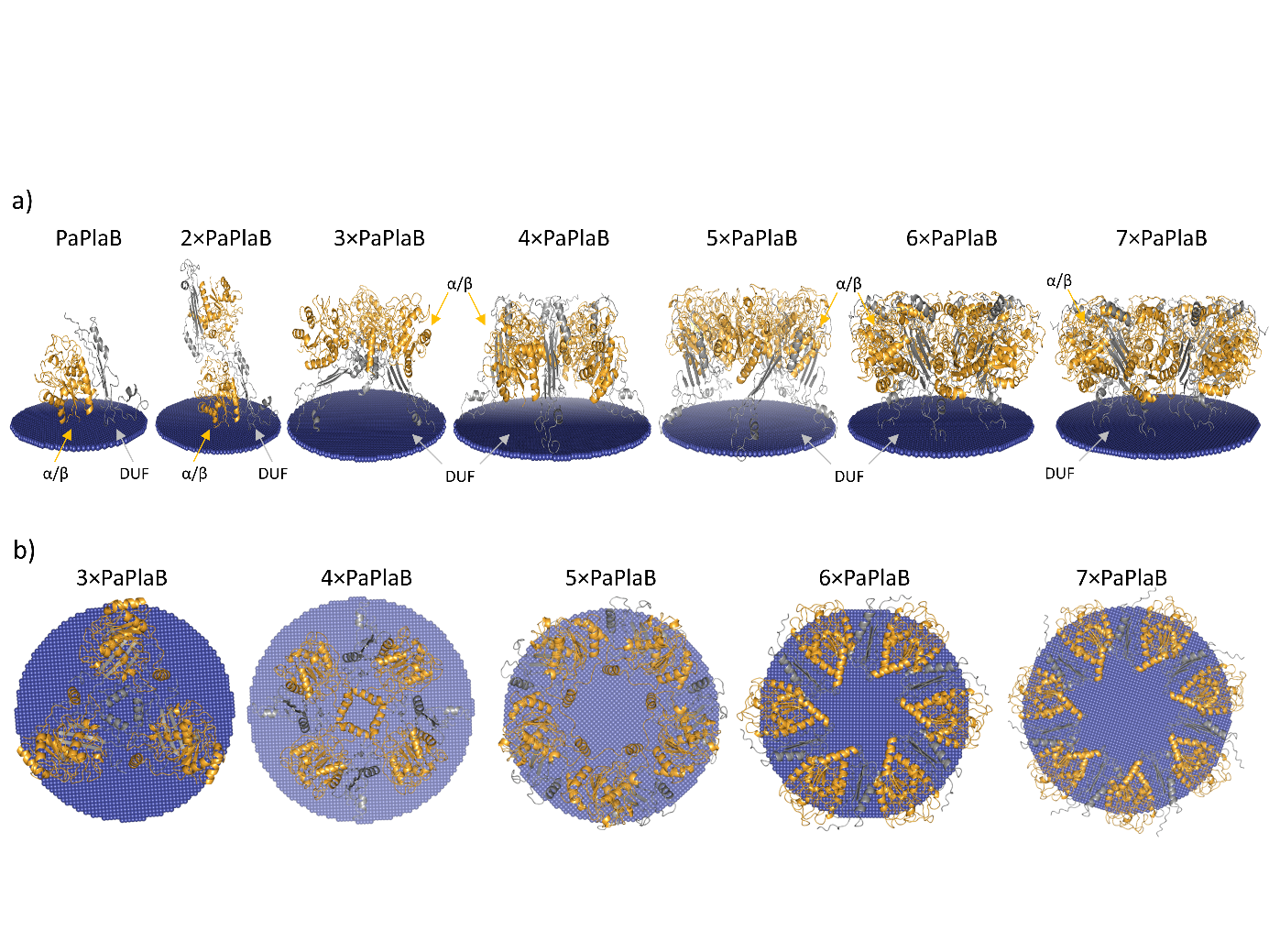

Fig. S10: Prediction of PaPlaB oligomer structures. a) PaPlaB monomer, shown in figure 1b, dimer (2xPaPlaB), and higher oligomers (3x – 7xPaPlaB), which were oriented at the membrane (blue). The catalytic α/β-hydrolase domain and DUF are colored yellow and grey, respectively. b) The ring-like structure of 3x – 7xPaPlaB observed from the top on the membrane.

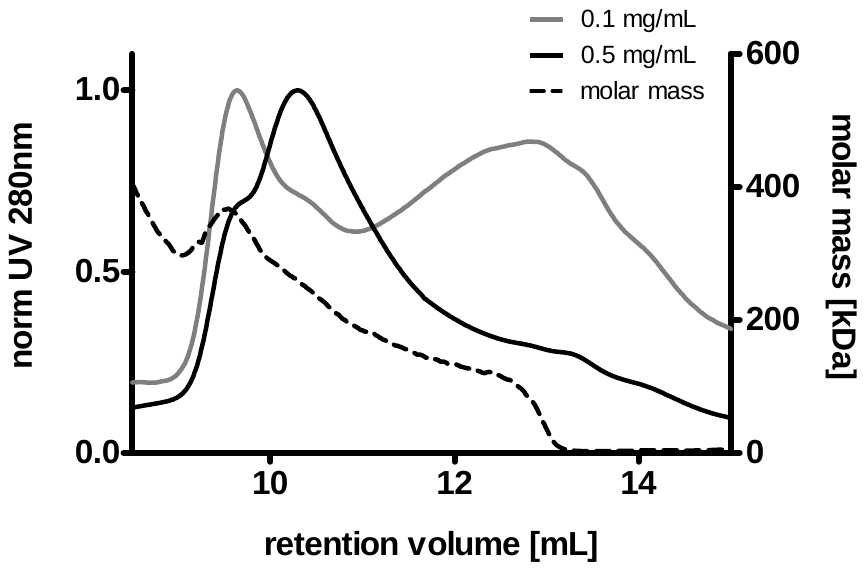

### Fig. S11: SEC-MALS analysis of PaPlaB at 0.5 mg/ml (black line) and 0.1 mg/ml (grey line). The molar mass (dashed line) was determined using ASTRA 7 for 0.5 mg/ml only. For 0.1 mg/ml no reliable mass determination was possible since the signal in the refractometer was too low.

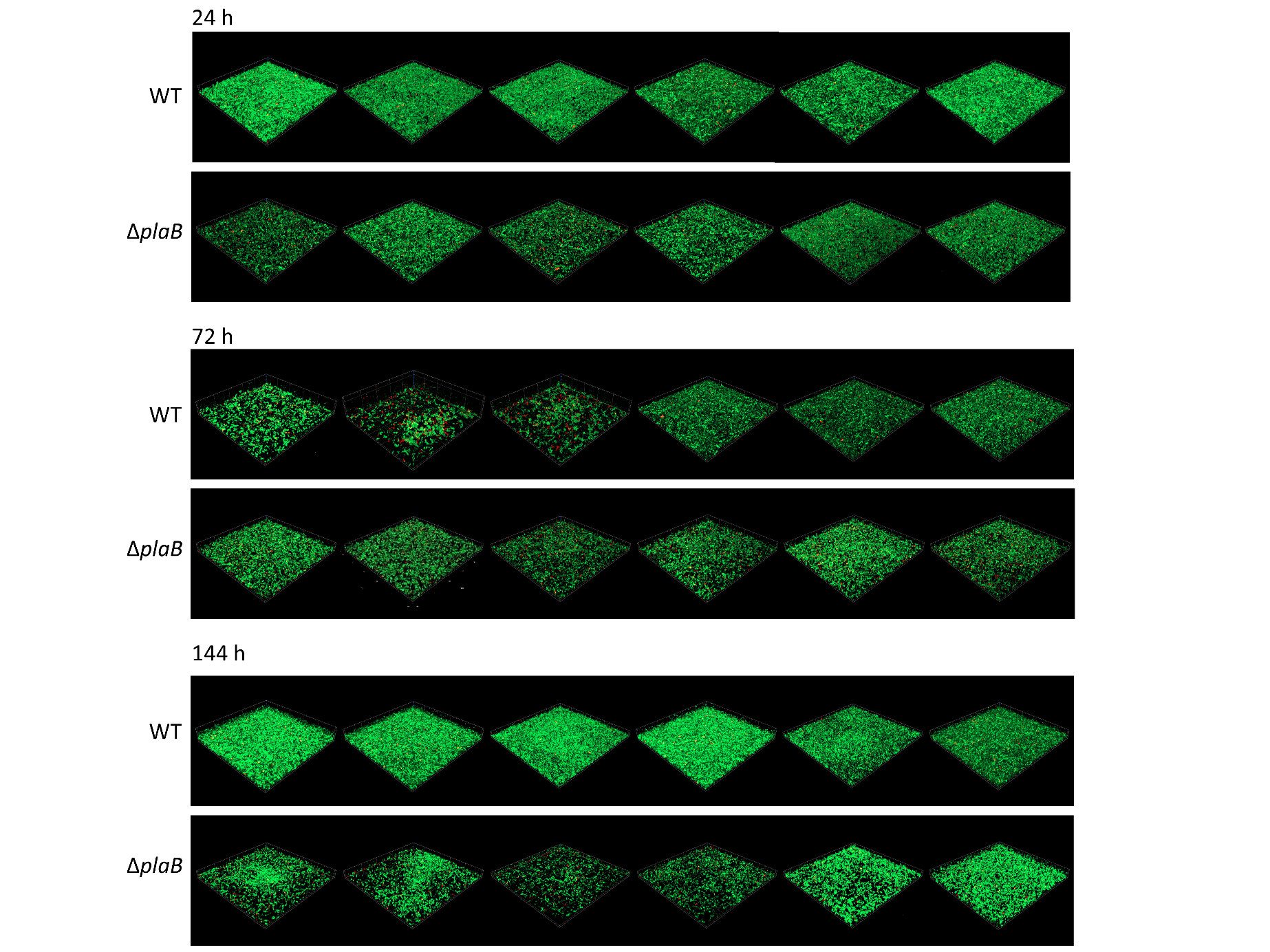

**Fig. S12: CLSM analysis of biofilm architecture.** *P. aeruginosa* wild-type (WT) and Δ*plaB* were grown at 37ºC in flow cell chambers with continuous supply (50 µl/min) of LB medium for 24, 72, and 144 h. Results were obtained from two biological replicates by analyzing biofilm at three different points. Figures show sections with a size of 100 x 100 µm.
